# Drug-like small molecules that inhibit expression of the oncogenic microRNA-21

**DOI:** 10.1101/2022.04.30.490150

**Authors:** Matthew D. Shortridge, Bhawna Chaubey, Huanyu J. Zhang, Thomas Pavelitz, Gregory L. Olsen, George A. Calin, Gabriele Varani

## Abstract

We report the discovery of a series of drug-like small molecules which bind specifically to the precursor of the oncogenic and pro-fibrotic microRNA-21 with mid-nanomolar affinity. These molecules are highly ligand-efficient (MW<330) and display specific biochemical and cellular activity by suppressing maturation of miR-21, thereby providing an avenue towards therapeutic intervention in multiple diseases where miR-21 is abnormally expressed. The small molecules target a local structure at the Dicer cleavage site and induce distinctive structural changes in the RNA which correlate with specific inhibition of miRNA processing. Structurally conservative single nucleotide substitutions eliminate the conformational change, which is not observed in other miRNA precursors. The most potent of these compounds reduces cellular proliferation and miR-21 levels in cancer cell lines without inhibiting kinases or classical receptors, while closely related compounds without this specific binding activity are inactive in cells.

## Introduction

Human microRNA-21 (miR-21) is a well-known proto-oncogene, marker of fibrosis and molecular link between inflammation and various disorders, including irritable bowel disease ^1-9^. When upregulated in response to inflammatory signals, the mature miR-21 post-transcriptionally silences hundreds of genes that regulate multiple biological pathways, including multiple tumor suppressors (e.g., PDCD4, PTEN), acting in a pleiotropic fashion ^10-12^. In multiple mouse models, miR-21 plays a causal role in malignant transformation, metastatic spread or resistance to treatment, suggesting that pharmacological inhibition of this ‘oncomiR’ could reverse even late-stage cancer ^1, 2, 13, 14^.

The mature miR-21 is highly conserved in sequence and generated by the canonical microRNA biogenesis pathway. The functional mature miR21-5p sequence, located on the 5′-strand of pre-miR-21, between residues U8 and A29 is overexpressed in many disease conditions (Fig. S1; as numbered from the primary sequence in miRbase). The direct targeting of mature miRNA sequences using antisense oligonucleotide chemistries has been reported in numerous studies, with encouraging results in cellular and animal models but disappointing clinical outcomes and significant toxicities ^2, 4, 13, 15, 16^. Because of toxicities, and the well-known limitations in delivery and distribution ^16^, there remains a compelling need to discover small molecules that inhibit the biogenesis pathway that generates the pathogenic mature miR-21, thereby reducing its over-expression in disease.

Many small molecules have been reported in the literature to bind to pre-miR-21, regulate its expression ^17^ and generate phenotypic responses ^18^, but the evidence of direct cellular engagement has been universally weak. For example, natural products discovered in an enzymatic screen are unlikely to be specific^19^, resulting in activity that is independent of RNA binding ^20, 21^. Mitoxantrone ^22^ is a cytotoxic PAIN molecule ^23^ that binds to many RNAs ^24, 25^; the NMR spectrum of its complex with pre-miR-21 is however extensively broadened due to the non-specific nature of the interaction (Fig S2). Peptoids are too flexible to bind specifically ^26-28^, while peptides selected by phage display were insoluble in our hands ^29^. Our 14-mer macrocyclic peptides had only weak inhibitory activity in biochemical assays, indicative of a superficial interaction (Fig. S2) ^30^; although we were able to improve affinity and especially biochemical potency by using non canonical side chains ^31^ (manuscript in preparation), peptides present numerous pharmacological challenges. In targeting other unrelated microRNAs, sub-uM affinity was reported only with molecules which do not satisfy the ‘rule of 5’ criteria of successful pharmaceutical projects ^32, 33^, making them useful only as probes. Finally, Ribotac strategies have been proposed ^34, 35^, but the pharmacological potential of these molecules remains to be demonstrated. Altogether, the identification of pharmaceutically attractive small molecules which bind to miRNA precursors potently and specifically yet retain drug-like properties remains elusive.

In investigating a class of RNA-binding small molecules that bind to HIV TAR RNA (BIORXIV/2022/477126), we discovered that compounds incorporating a 2-((5-(piperazin-1-yl)pyridin-2-yl)amino)pyrido[3,4-d]pyrimidin-4(3H)-one structure with the carbonyl functional group situated within the pyrimidine fragment (Fig. 1 and Table 1), display strong binding activity to pre-miR-21 in biophysical assays and inhibit processing of the precursors by the enzymes that generate mature miR-21 in rigorous biochemical assays and in cells. These Lipinski molecules are very ligand-efficient (MW about 330) and target a specific structure at the junction between the apical loop and helical stem of pre-miR-21 which is required for processing by both Drosha and Dicer. This affinity is reduced significantly by conservative single nucleotide changes in the pre-miRNA. The most potent compounds derived from this structure have binding activity into the mid-nM range, and simple modifications of the structure, such as moving the position of a nitrogen within a ring, reduce binding significantly, demonstrating clear structure-activity relationships; compounds within the series which lack the same binding profile are inactive in cells.

**Table 1.**
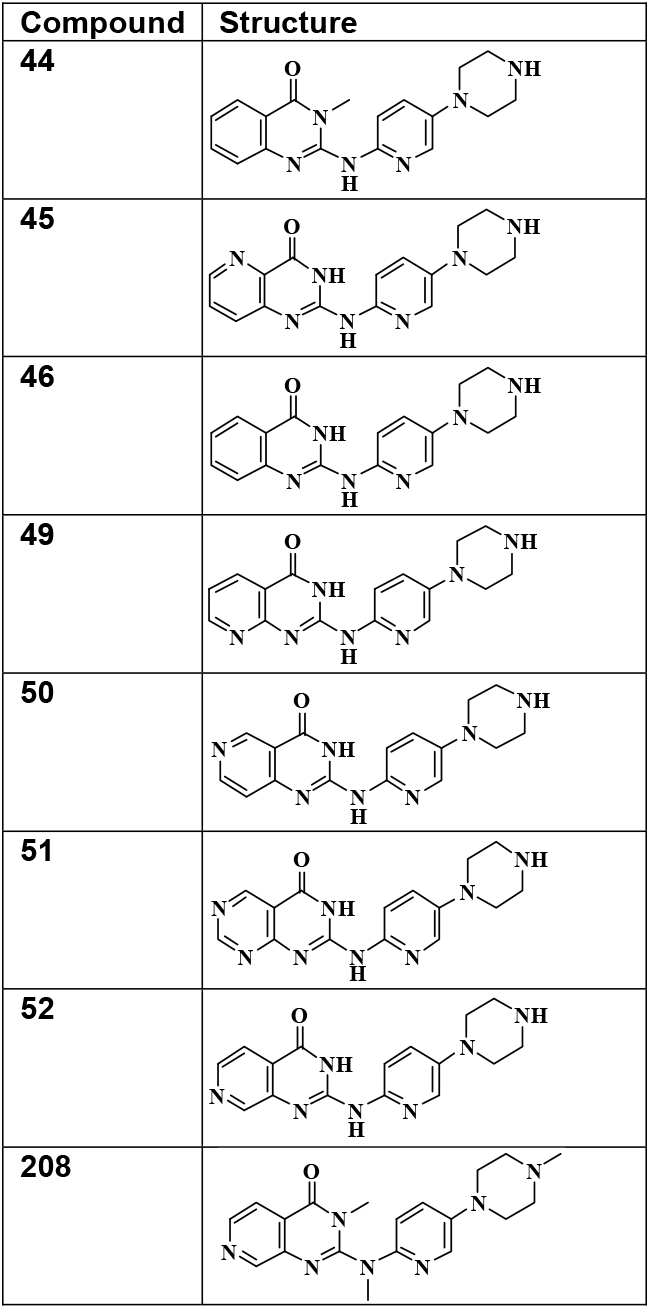
Structure of compounds described in the manuscript.

**Figure 1.**
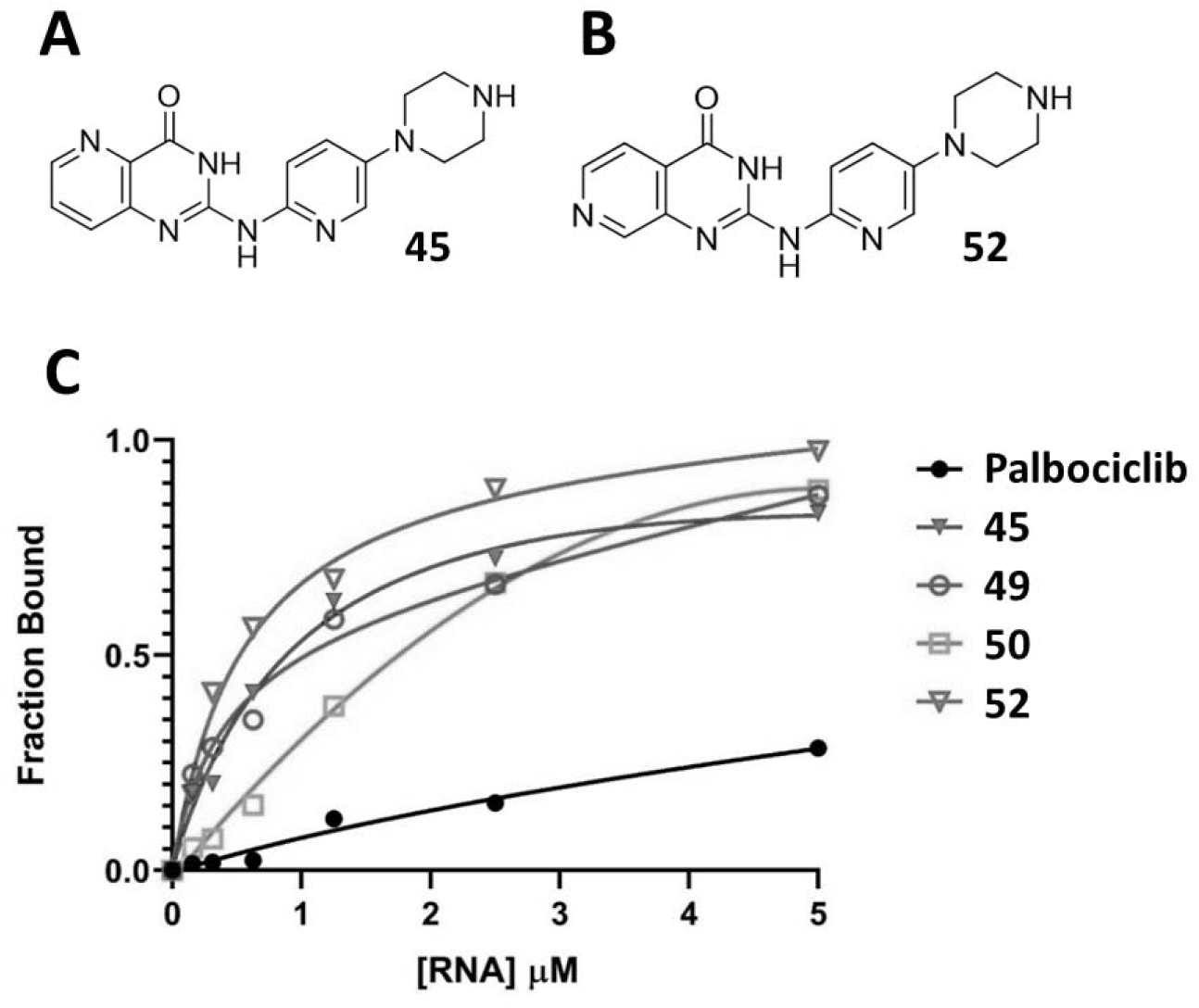
Drug-like small molecules bind to pre-miR21. We synthesized a series of closely related compounds based on a common 2-((5-(piperazin-1-yl)pyridin-2-yl)amino)pyrido[3,4-d]pyrimidin-4(3H)-one structure and assayed them for binding to pre-miR-21 using a universal NMR assay ^1, 2^. Two compounds, called 45 (A) and 52 (B), had strong binding activity, in the mid-nM range. Compounds generated by moving the position of a single nitrogen (Table 1), display significantly reduced affinity (5-10-fold differences) (C). 1D ^1^H NMR ligand detected titrations were performed to assess binding of the candidate compounds: increasing concentrations of RNA were added into a solution containing 100 LM of the small molecule in a buffer containing 50 mM deuterated Tris at pH 6.5, as well as 250 mM NaCl, 50 mM KCl and 2 mM MgCl_2_. As increasing amounts of small molecules bind to the RNA, the ^1^H linewidth increases, and the NMR peak height correspondingly decreases. The fraction of bound small molecule is calculated from the decreases in peak height relative to an internal standard (DSA). Saturation of the curves to a value of 1 indicates the presence of a dominant single site with sub-uM affinity; by comparison, an unrelated RNA-binding compound, Palbociclib, saturates at a much lower value and shows a nearly linear titration curve, indicative of non-specific binding (see Table 1 for structures of all compounds tested). Approximate binding constants can be calculated by fitting the data points to a binding isotherm. The data fit for compound 52 corresponds to an approximate K_d_ = 200 nM, while compounds 45 and 49 (Table 1) both have K_d_ = 600 nM.

Reducing cellular levels of miR-21 with small molecules targeting miRNA biogenesis is expected to have therapeutic benefits in a variety of cancers ^1, 2, 14, 35^ as well as fibrotic and inflammatory conditions ^4-6, 36^. The small molecules described herein, by residing in the conventional favorable ‘Lipinski’ drug-like chemical space ^37^, thus hold considerable promise to address an unmet need in RNA-targeted pharmacology.

## Results

### The Dicer binding site displays a different conformation in different pre-miR-21 constructs

Most NMR work on pre-miR-21 has studied short RNA stem-loop models ^30, 38^, as shown in Figure S1. We reported the three-dimensional structure of this RNA alone and in complex with a small cyclic peptide (PDB 5UTZ and 5UZZ). In that construct, the A29 nucleotide at the 5’ Dicer cleavage site is stacked within the RNA helix, and we observe clear nuclear Overhauser effect (NOE) interactions between the amino nitrogen protons (NH_2_) of A29 and the imino resonances of both G28 (11.6ppm) and G45 (12.5ppm). The A29 NH_2_ resonances ^38^ resonate at 8.9ppm and 10.1ppm, consistent with formation of a protonated A^+^-G A29:G45 base pair. However, the full pre-miR-21 is a 59 nt hairpin, and the characteristic NOE pattern arising from A29 in the short construct is not observed in the full pre-miR-21 (BIORXIV/2021/471640; the upfield-shifted A29 NH_2_ signal can be seen in Fig. S1B only for the short construct), demonstrating that the two sequences have different structures at the helical-loop junction region — i.e. the Dicer cleavage site. In this study, we use either the full length pre-miR-21, or an intermediate length sequence (Fig. S1) that recapitulates the structural and dynamic properties of the full pre-miR-21 which are missing in the shorter stem-loop model. All sequences studied are listed in Table S1.

### Drug-like small molecules bind to pre-miR-21 with mid-nM affinity and induce a new structure near the Dicer cleavage site

We recently reported that Palbociclib binds with high affinity to HIV TAR RNA and designed a variant molecule that retains binding activity but does not inhibit kinases (BIORXIV/2022/477126). Because that molecule was toxic, we synthesized additional variants to eliminate the toxicity, using pre-miR-21 as a control for specificity during this elaboration. We discovered that a 2-((5-(piperazin-1-yl)pyridin-2-yl)amino)pyrido[3,4-d]pyrimidin-4(3H)-one structure, with the carbonyl functional group positioned within the pyrimidine fragment, instead provided strong binding activity to pre-miR-21, while binding to TAR was significantly decreased (Fig. S3).

Based on that unexpected result, we synthesized a series of analogues (Table 1), and used a ligand-detected NMR relaxation assay (Fig. S3 and S4) to obtain their approximate binding affinities. Among the small number of compounds synthesized, compounds 45 and 52 showed Kds of approximately 600 nM and 200 nM, respectively (Fig. 1 and S3); most significantly, small structural modifications to these compounds, such as shifts in the position of a nitrogen, or removal or addition of a nitrogen to a 6-membered ring (see Table 1), resulted in 5-10 fold losses in binding activity (compare compound 52 with compounds 45, 49, and 50, which differ only in the positioning of a nitrogen within the six-membered ring; compounds 45 and 49 are about 3-fold weaker binders, while the very similar compound 50 has significantly weaker binding activity). Such pronounced changes in response to fine-tuning of local chemical details are rarely observed among RNA-binding small molecules, where SAR is often shallow and relatively small changes in affinity are instead seen even in response to much more radical chemical changes ^39, 40^.

The strong binding activity of compounds 45 and 52 is correlated with distinctive structural changes in the RNA. First, each ‘closes the loop’ of pre-miR-21 (Figs. 2A and 2C). Namely, titration of pre-miR-21 with compounds 45 and 52 results in the emergence of new imino peaks assigned to the apical loop, corresponding to formation of two GU base pairs just above the Dicer cleavage site. Further confirmation that these were GU base pairs was obtained by recording ^15^N-edited HSQC spectra (Fig. S5).

**Figure 2.**
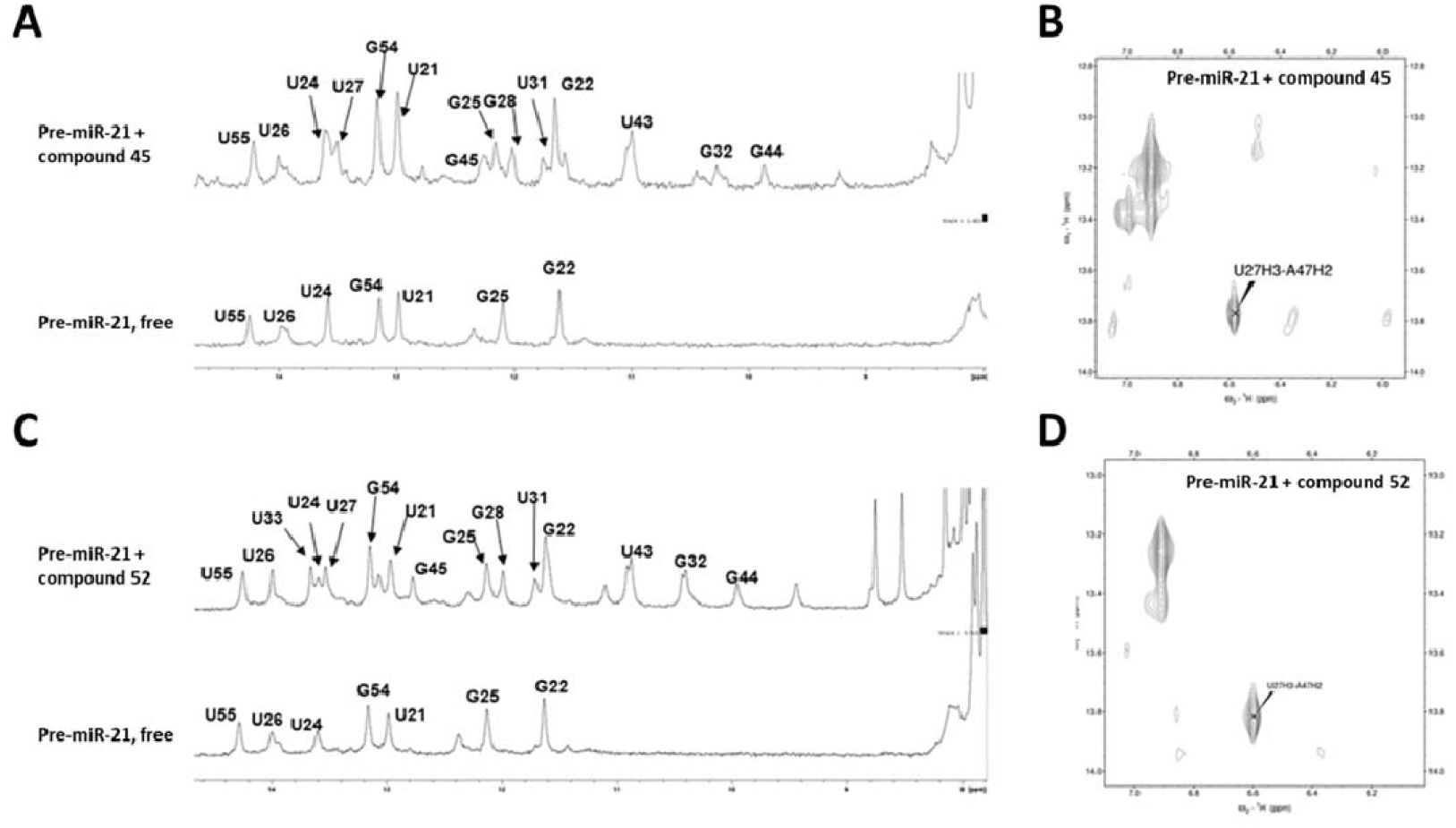
1D ^1^H NMR and NOESY spectra for pre-miR-21 with and without addition of compounds 45 and 52. Compounds that bind strongly to pre-miR-21 (45 and 52) ‘close’ the apical loop of pre-miR-21 and stabilize a conformation of pre-miR-21 that is processed inefficiently by Dicer-TRBP. Compound 45 (**A, B**) and compound 52 (**C, D**) are unique among numerous pre-miR-21 binding compounds we have investigated. The 1D ^1^H NMR spectra focused on the imino region (**A** and **C**), reveal closing of the two GU wobble base pairs in the apical loop. (**A**) A 1mM sample of pre-miR-21 was titrated to 1.5mM with compound 45; new resonance signals appearing for U31, G32, U43 and G44 (A, top) demonstrate formation of the tandem UG/GU wobble pair in the apical loop (Fig. S5), but the peaks are split, indicating the presence of two nearly equally populated conformations. Compound 52 (**C**) induces a similar but stronger response under the same conditions, closing the U33:A42 base pair as well, in addition to the two new GU pairs. Both compounds also select a unique conformational state of the pre-miR-21 hairpin as demonstrated by NOESY spectra recorded under the same conditions (**B** and **D**): the U27H3-A47H2 NOESY signal labeled in each spectrum shows the close contact between U2)7 and A47 within that base pair and corresponds to the inefficiently processed A29-bulged out conformation of the Dicer cleavage site (**BIORXIV/2021/471640**).

A second structural consequence is the stabilization of a bulged-out conformation of A29 that corresponds to a state that Dicer processes inefficiently (BIORXIV/2021/471640). In free pre-miR-21, A29 occupies two nearly equally populated conformations. This is manifested as a splitting of the U27 NH and A47 H2 NOE resonances, corresponding either to stacking of the base within the helix or unstacking and repositioning outside of it (Fig 2B and 2D). Binding of 45 or 52 strongly enhances the downfield shifted U27 NH signal arising from the bulged-out A29 state, a conformation which represses pre-miR-21 processing (BIORXIV/2021/471640).

### The interaction between the small molecules and pre-miR-21 is highly specific

To evaluate specificity, we tested binding of compound 52 to HIV TAR, and observed a >10-fold decrease in affinity compared to pre-miR-21 (Fig. S3B). In contrast, two other high-affinity TAR ligands that we have recently reported bind 20-30 less potently to pre-miR-21 than to TAR (BIORXIV/2022/477126).

To more surgically assess specificity, we introduced conservative mutations at or near the sites in pre-miR-21 where the conformational change takes place (Table S1 and Fig. S6A). Remarkably, inversion of the U33-A42 base pair to A-U reduces binding by about 5-10-fold (Fig. 3, mutant 3). Even more dramatic effects are observed when the GU base pair adjacent to U33-A42 is changed from GU to GC (mutant 1), or the base pair just below is also changed from UG to CG (mutant 2) (Fig. 3 and S6). These are structurally conservative substitutions; introduction of changes more radical than these is not uncommon when testing for specificity, for example the removal of the unpaired A29 to generate a perfect helix that is refractory to binding small molecules.

**Figure 3.**
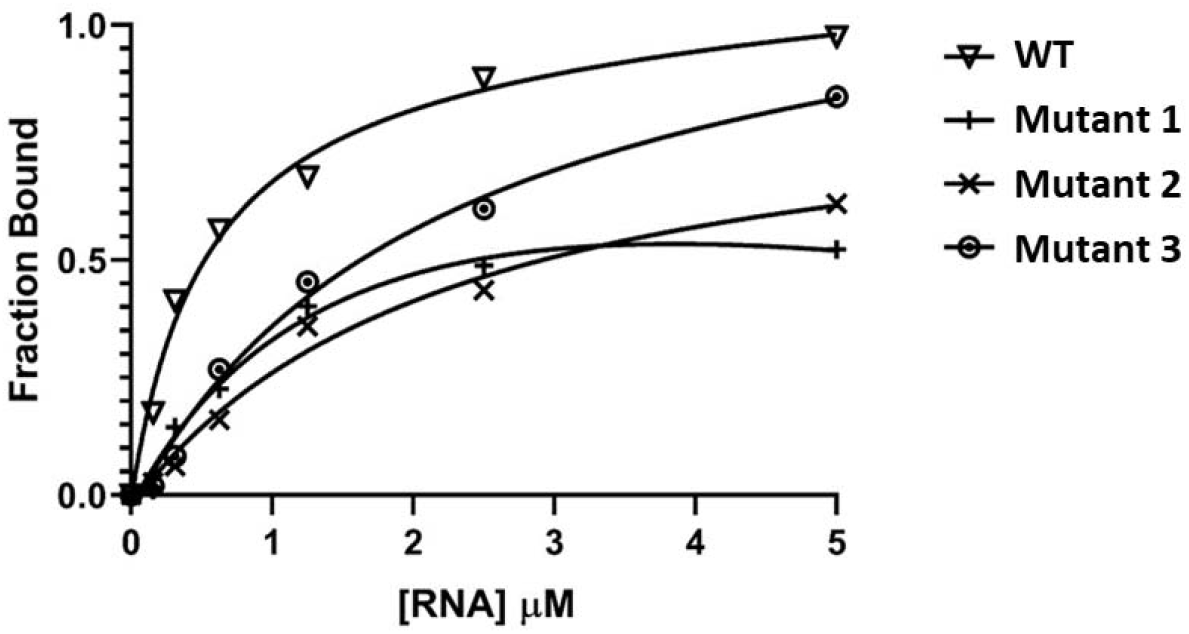
Ligand-detected NMR titration of compound 52 with pre-miR-21 and three mutant RNAs. Binding of compound 52 to pre-miR-21 is sensitive to small, conservative single- or two-nucleotide substitutions. Changing single base pairs from GU to GC (mutants 1 and 2), or inversion of an AU base pair to UA (mutant 3) leads to 5-10-fold reductions in affinity compared to wild type pre-miR-21 (inverse triangle), as demonstrated in ligand-detected NMR titration experiments. (See Fig. S6 for RNA sequences).

An explanation for these relatively large effects is provided by the analysis of the NMR spectra; once the double mutation is introduced to invert the UA base pair, the closed-loop conformation induced in the wild-type pre-miR-21 by compound 52 is destabilized and no longer observed (Fig. S6B).

To further test for specificity, we challenged several other pre-microRNAs (Table S2) with molecules 45 and 52. With pre-miR-24, pre-miR125b, and pre-miR10b, we observe only small changes in the NMR spectra upon binding of stoichiometric amounts of 45 or 52, further demonstrating that the interaction is specific for pre-miR-21 (Fig. S7). This is particularly noteworthy for pre-miR-10b, which shares the same UGU element recognized by the DGCR8 component of the microprocessor complex ^41^. Notably, we do not observe stabilization of the base pairs in pre-miR-10b, and very few intermolecular NOEs (Fig. S8A) are present in NOESY spectra. For pre-miR-21, in contrast, numerous intermolecular NOEs are seen between the RNA and the small molecule (>50, Fig. S8B), consistent with a high affinity and site-specific interaction. A preliminary analysis of the pre-miR-21 NOESY data (Fig. S5) identifies close NOE contacts between protons on the compound 52 heterocycle (two distinctive peaks at 8.5-9 ppm) and the NHs of the two consecutive GU base pairs, consistent with the formation of direct intermolecular contacts between the small molecule and the major groove edges of the GU base pairs, resulting in the stabilization of the bound structure.

### The small molecules that bind to pre-miR-21 have specific biochemical activity

The NMR analysis demonstrates that molecules 45 and 52 ‘close the loop’ by stabilizing the GU base pairs, and at the same time induce the A29-out conformation that represses miR-21 processing by Dicer (BIORXIV/2021/471640). Encouraged by this observation, we evaluated their biochemical activity using both a commercial enzyme preparation and a highly active recombinant Dicer-TRBP preparation expressed *in house*.

First, we titrated pre-miR-21, with increasing amounts of compound 52, or of an inactive variant (compound 208) generated by methylating the three NH functionalities of the molecule. This second molecule does not bind to pre-miR-21 even at 10 uM concentration (Fig. S9) and does not induce inhibition of pre-miR-21 processing even at concentrations >50 uM. In contrast, molecule 52 inhibits pre-miR-21 processing by Dicer with a single digit uM approximate inhibition constant (Fig. 4 and S10).

**Figure 4.**
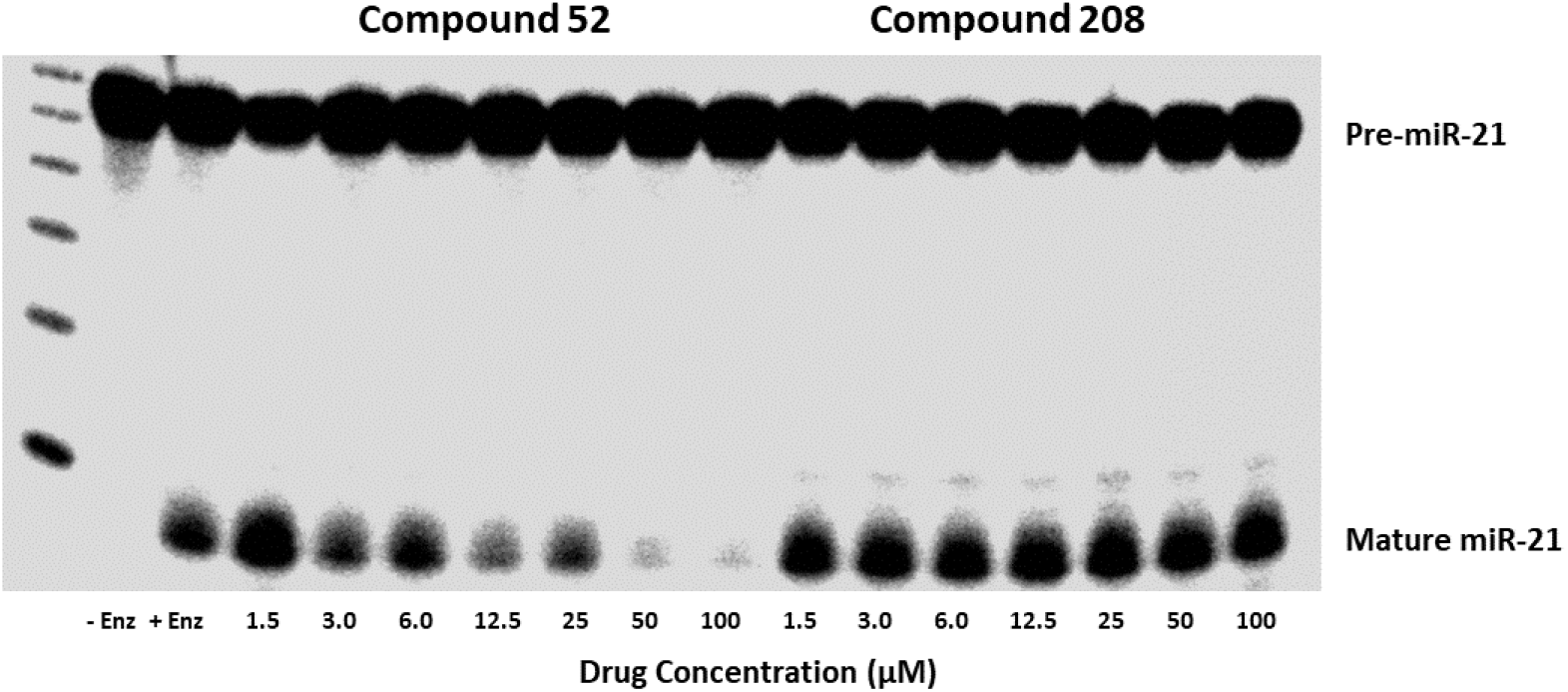
Pre-miR-21 processing by Dicer in the presence of increasing amounts of compounds 52 and 208. Inhibition of pre-miR-21 processing by Dicer is observed after addition of compound 52, but not for a version of the same compound that has lost binding activity through methylation of NH functionalities (Fig. S9). Processing reactions were run for 1 hour in the presence of various concentrations (1.5, 3, 6, 12.5, 25, 50, and 100 µM) of either compound 52, or compound 208, and an aliquot run on a 15% sequencing gel. The leftmost lane contains size markers from a 10bp DNA ladder.

The in house-purified Dicer-TRBP processes pre-Let-7a and pre-miR-21 in minutes, similar to processing times within the cell (Fig. S10), which allows inspection of the early (minutes) time points in the reaction, before significant product builds up. With pre-Let7a, no significant difference in activity was observed in the presence or absence of 2 uM of compound 45. In contrast, at this same concentration, compound 45 significantly reduced pre-miR-21 processing by Dicer-TRBP, relative to no compound control.

### Pre-miR-21 binding compounds have specific cellular activity

Three sets of measurements were conducted in cell lines to evaluate cellular responses after exposure to the pre-miR-21 binding compounds: (i) proliferation of transformed cell lines, (ii) levels of mature miR-21, and (iii) levels of the downstream target PDCD4. All assays were conducted in a blind format at the MD Anderson cancer center, without knowledge of the identities or binding activities of the compounds tested.

The anti-proliferative activity of compounds 45 and 52 was determined in two cancer cell lines characterized by high levels of miR-21, gastric adenocarcinoma AGS, and pancreatic ASPC1. Compound 52 and Palbociclib, a positive control as a well-known cancer inhibitory drug in both AGS ^42^ and ASPC1 cells ^43^, both showed anti-proliferative activity at both concentrations tested in standard MTS assays conducted over 6 days. Cellular viability in the presence of either 52 or Palbociclib at 10 (Fig. 5) or 5 nmol/ml (Fig. S11) decreased significantly after approximately 72-96 hours of culture, but no significant effect was observed for compound 45.

**Figure 5.**
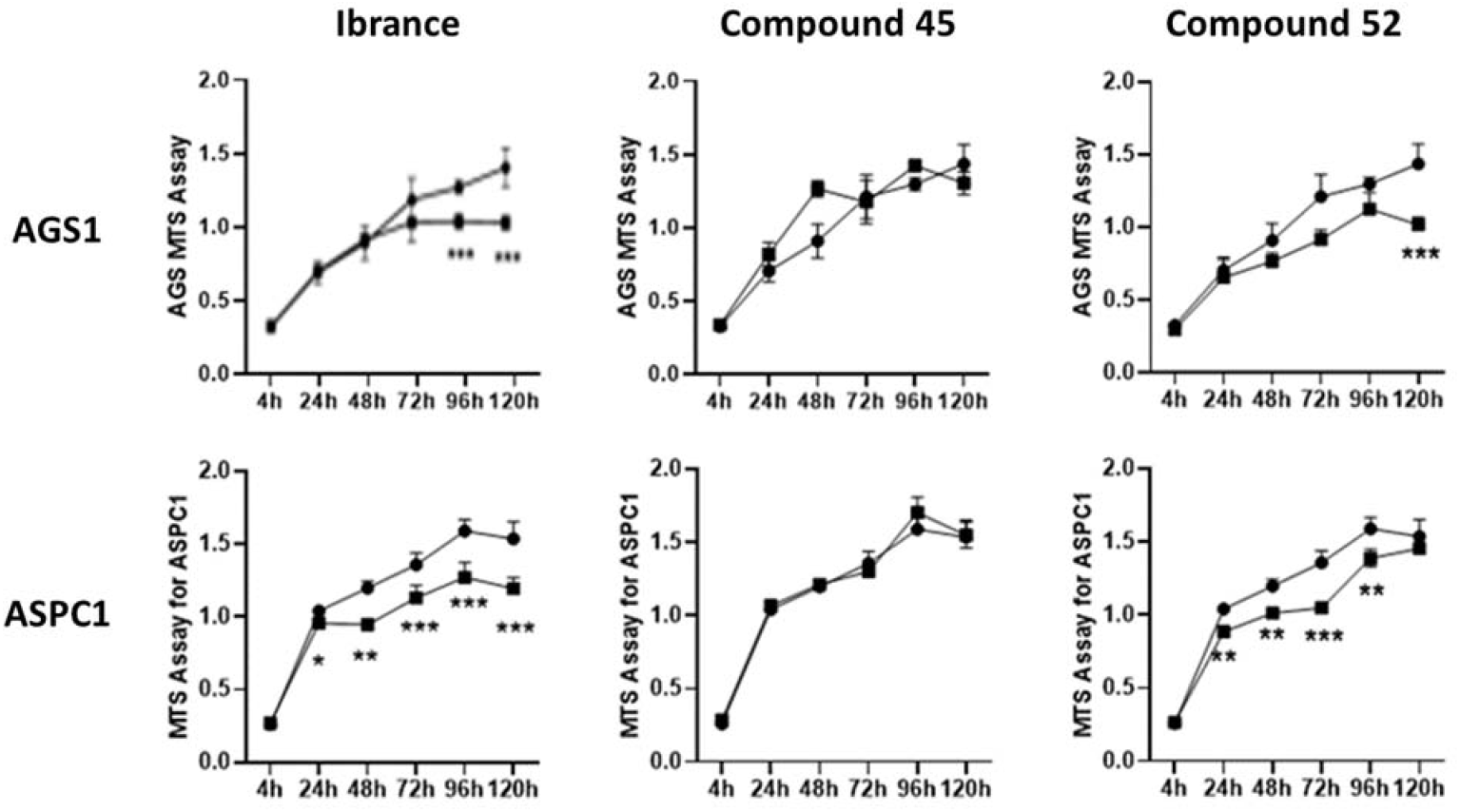
Anti-proliferative activity of compounds 45 and 52 against gastric (AGS) and pancreatic (ASPC1) cancer cell lines, measured as a function of time in a blind experimental format and compared to the breast cancer drug Palbociclib (Ibrance). Cell viability was assayed using standard MTS assays in the indicated cell lines, following addition of each compound at 10 uM concentration (10 nmoles/ml); error bars show experimental uncertainty from results collected in triplicate. Palbociclib and 52 reduce the viability of both cell lines to a statistically significant extent (marked with asterisks) while 45 does not have any significant effect.

Levels of mature miR-21 and its precursor pre-miR-21 were measured in the same cancer cell lines under conditions where we observed a decrease in proliferation (Fig. 6). Significant (30%) decreases were observed in mature miR-21 levels in both AGS and ASPC1 cells when challenged with 45 or 52, while inconsistent changes were observed for the control Palbociclib, a kinase inhibitor not expected to reduce miR-21 levels (Fig. 6). This level of reduction is as much as can be expected for an inhibitor of microRNA maturation, because microRNAs are long lived (2-5 days) ^44^ and only newly produced mature miR-21 would be affected by our compounds. Large reductions of mature miR-21 levels over short time scales (24-48 hrs), as reported ^18, 22^, would not be consistent with inhibition of new microRNA expression. Reductions in mature miR-21 were also accompanied by a reduction in pre-miR-21 levels, suggesting that at least part of the inhibitory block occurs prior to Dicer processing.

**Figure 6.**
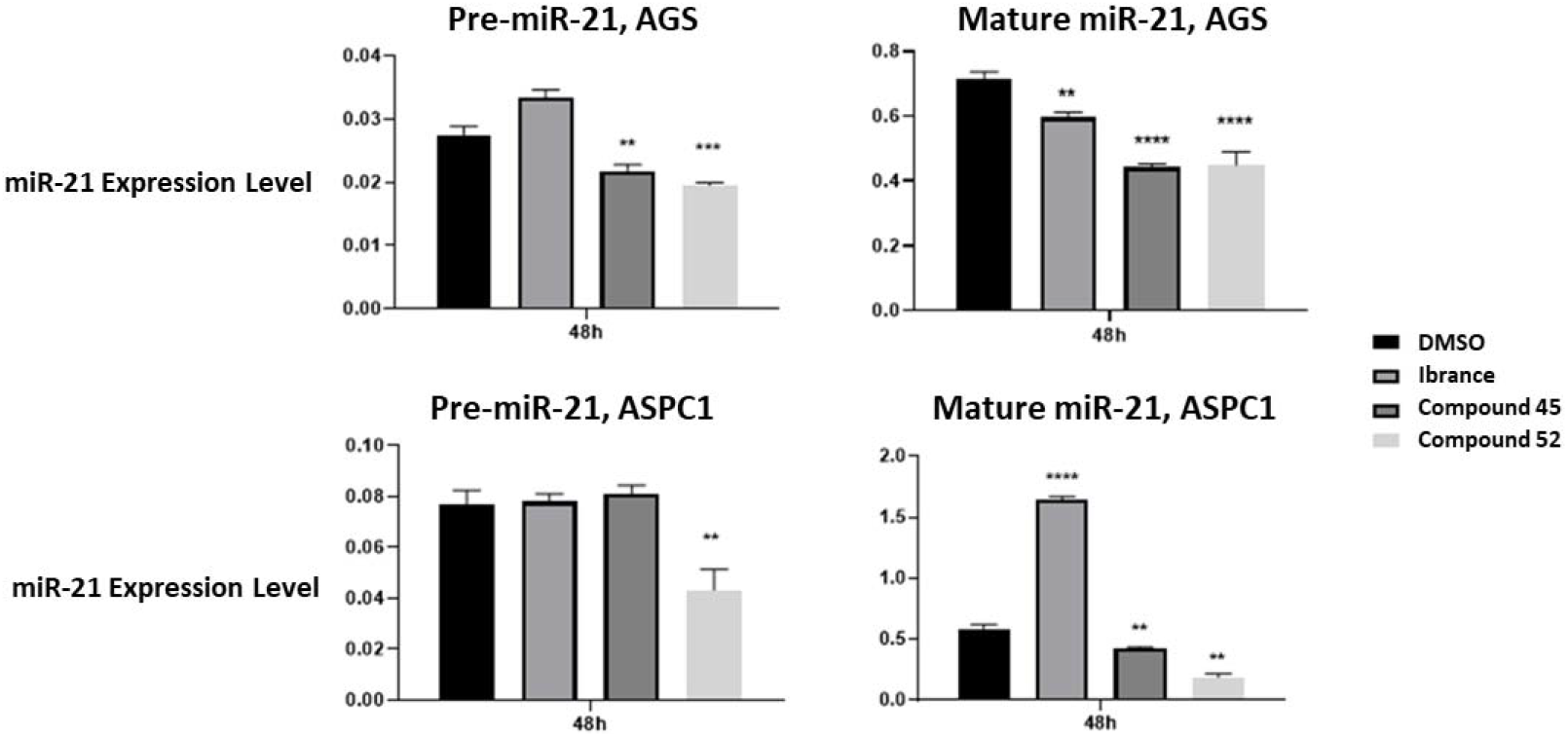
Levels of pre-miR-21 (left) and mature miR-21 (right) measured in gastric adenocarcinoma (AGS) and pancreatic cancer cell lines (ASPC 1), normalized to internal controls (U6 snRNA), in the presence of 10 nmoles of Palbociclib (used here as a negative control not expected to affect microRNA levels) and of compounds 45 and 52, measured 48 hours after incubation, relative to DMSO control, and measured by standard qRT-PCR. Statistically significant changes are labeled with asterisks.

As shown in Fig. S12, inhibition was specific to miR-21. Levels of mature miR-16 ^45^ (a tumor suppressor miRNA), miR-28 ^46^ (an oncogenic miRNA in intestinal cancers) and miR-182 ^47^ (an oncogenic miRNA, whose expression is closely related to the clinicopathological features of gastric cancer) are not consistently or significantly affected by the compounds in either gastric (AGS) or pancreatic (ASPC1) cell lines, over multiple time scales (24-72 hours), relative to controls where only DMSO was added.

In order to assess the molecular mechanism of inhibition of miR-21 expression, we also investigated some of the lower-affinity compounds in the series (44, 46, 50 and 51, Table 1). In both AGS and ASPC1 cells lines (Fig. S13), we observe no significant changes in miR-21 levels upon addition of these compounds, which bind to pre-miR-21 less potently than compounds 52 or 45. Furthermore, levels of both U6 and U8snRNA are not affected at all, demonstrating that our compounds do not affect general RNA metabolism.

Finally, when levels of PDCD4 (a known target of miR-21 ^11, 48, 49^) were measured, an increase in the levels of the tumor suppressor was induced by both compounds 45 and 52 after 48 hours of incubation (FIG. S14), showing that the decrease in mature miR-21 levels also leads to an increase in the PDCD4 downstream target of miR-21.

### Compounds 45 and 52 show only minimal kinase or receptor activity in panel screen assays

We conducted two additional experiments to investigate whether binding to protein targets could be responsible for the observed effects. First, we executed a KinomeEdge assay, measuring inhibition of the activity of approximately one hundred human kinases. Only very weak inhibition (>10 uM) of a small number of kinases was observed for both compounds 45 and 52, strongly suggesting that the effects we observe are not due to kinase inhibition (Fig. S15). Second, we conducted a CEREP assay, to measure activation or suppression of the activity of about 30 classical drug receptors (Fig. S16). Only very weak activation of the cannabinoid receptor was observed, with no other significant effect in agonizing or antagonizing any other receptor. This assay is generally used to flag potential toxicities early in drug discovery programs, but the results indicate no broad activity against classical receptors, in addition to a safe early pharmacological profile.

Taken together, these data indicate that not only does compound 52 induce decreased cell proliferation for AGS and ASPC1 cells, the same compound also displays RNA-specific responses in cells, decreasing levels of mature miR-21 and restoring levels of the tumor suppressor PDCD4, while other microRNAs are unaffected. Furthermore, inhibition of miR-21 processing by compounds 45 and 52 is specific (miR-16, -28 and -182 are not affected) and not simply a result of reduced proliferation (Palbociclib does not significantly affect miR-21 levels).

## Summary

We report the discovery of a small series of drug-like small molecules that satisfy all Lipinski rules of pharmacologically attractive small molecules ^37^ and bind to the precursor of the pro-oncogenic and pro-fibrotic miRNA-21 with mid-nM affinity and specificity, and with high ligand efficiency. These molecules are unlike previously reported ligands for pre-miR-21 or other microRNAs, which typically had much weaker affinity, were not Lipinski molecules, were cytotoxic or much larger than most successful small molecule drugs. The two most potent compounds induce specific changes in the structure and dynamics of pre-miR-21 at the Dicer cleavage site, locking the RNA into an inefficiently processed state. As a consequence, our compounds reduce processing by Dicer *in vitro* and in cells and do so without affecting other microRNAs or cellular RNAs. We observe concomitant decreases in levels of mature miR-21 in cell lines of gastric and pancreatic cancer origin after compound addition, as well as increases in the well-known target of miR-21, the tumor suppressor protein PDCD4. The two compounds do not inhibit kinases and have a clean profile against classical receptors, indicating that they reside in a pharmacologically-safe chemical space. The compounds are thus prime candidates for further therapeutic development in diseases where overexpression of miR-21 plays a causative role.

## Methods

### RNA Transcription

All RNAs for NMR or biochemical work were prepared *in house* using *in vitro* transcription on a large scale (typically 10mL) ^50, 51^ or synthesized by IDT for more efficient ^32^P labeling. RNA transcription and purification protocols used purified DNA oligonucleotide templates (IDT) and T7 RNA polymerase. Briefly, 1 mL of 8 uM DNA (5’-CTATAGTGAGTCGTATTA-3’), corresponding to the phage T7 RNA polymerase promoter region, was annealed to 80 uL of 100 uM template sequences with 13 mM MgCl_2_, heated to 95°C for 4 min then allowed to cool to room temperature over 20 min. Following annealing, the mixture was incubated in transcription buffer, 8% PEG-8000, and 35 mM magnesium chloride, with 5 mM of each of the four NTPs (ATP, GTP, UTP and CTP, from Sigma) and 0.4 mg/mL T7 RNA polymerase expressed and purified *in house*.

All RNA samples were purified from crude transcriptions by 20% denaturing polyacrylamide gel electrophoresis (PAGE), electroeluted, and concentrated by ethanol precipitation. The samples were re-dissolved in 12 mL of high salt wash (700 mM NaCl, 200 mM KCl, in 10 mM potassium phosphate at pH 6.5, with 10 µM EDTA to chelate any divalent ions), then concentrated using Centriprep conical concentrators (3,000 kDa MWC, Millipore). The RNA was then slowly exchanged into low salt storage buffer (10 mM potassium phosphate at pH 6.5, with 10mM NaCl and 10 µM EDTA). Prior to NMR experiments, all RNA samples were desalted using NAP-10 gravity columns, lyophilized and re-dissolved in buffer (see below), then annealed by heating for 4 min to 90 °C and snap cooling at -20 °C.

Sequences for the RNAs used in this study are shown in Tables S1 and S2.

#### Small molecule preparation

##### Sources

Palbociclib was purchased from Selleckchem. Starting materials were purchased from Absyn Chemicals (compound -52); or Combi-blocks, PharmaBlock, and Astatech.

##### General Procedure, Route-1 (Fig. S17)

The aniline (B, 1.05 equiv.) was taken up in dry 1,4-dioxane (0.1 M) in a microwave vial with a magnetic stir bar. 2-chloropyrimidine (A, 1.0 equiv.) was added followed by K_2_CO_3_ (3.0 equiv.) and X-Phos (0.2 equiv.). The reaction mixture (RM) was sparged with nitrogen gas for 5-10 minutes. Finally, Pd_2_(dba)_3_ (0.1 equiv.) was added, and the RM was sparged with nitrogen for another 5 min, after which the microwave vial was sealed, and microwave irradiated at 120 °C for 1.5 h. The RM was cooled to ambient temperature, filtered through a pad of Celite, rinsed with ethyl acetate and the solvent was removed under reduced pressure. The crude RM was purified on silica gel using 0-80% ethyl acetate in hexanes as eluent. Relevant pure fractions were evaporated in vacuum to give intermediate-C. Int-C was dissolved in DCM/TFA (4:1, 0.05 M), and stirred for 1-2 hours at RT. The crude reaction was concentrated and purified on HPLC using water/acetonitrile as eluent. Relevant pure peak fractions were lyophilized to generate compound -52 and analogs (overall yield for the two-step reactions: 20-50%).

##### General Procedure, Route 2 (Fig. S17)

1-Boc-4-(4-aminophenyl)piperazine (1.05 equiv.) and the corresponding 2-chloropyrimidine (A, 1.0 equiv.), TFA (3.0 equiv.) were taken in a sealed tube with n-BuOH (0.05 M). The RM was heated at 145 °C overnight (16-22 h). The RM was cooled down to ambient temperature and excess TFA was quenched with triethylamine (TEA). The crude compound was purified on HPLC using water/acetonitrile as eluent. Relevant pure peak fractions were lyophilized to give the corresponding -52 analogs (yield 35%).

### Ligand-detected NMR binding assay

Compounds were first dissolved to 1-10 mM in pure water or DMSO, depending on their solubility, then prepared to 100 uM in 490uL in 50mM bis-Tris, pH 6.5, in 50mM d_9_-deuterated bis-tris buffer at pH 6.5, containing 11.1uM sodium 4,4-dimethyl-4-silapentane-1-sulfonate (DSA) as chemical shift reference, all dissolved in 99.99% D_2_O. The non-binding internal DSA reference allows normalization of the small molecule spectra (in the absence or presence of RNA) during ligand-detected titrations. The 9 protons on the internal reference also provide a control for ligand concentration and flag compounds that aggregate or precipitate. All ligand-detected experiments were conducted using the 1D-^1^H NMR excitation sculpting water suppression scheme (Bruker sequence ‘zgesgp’); a free ligand reference spectrum was collected followed by titrations from stock RNA solutions up to 10uM RNA concentration in the NMR tube at the end of the titration. Each experiment was collected with 16 scans, 16k data points with a recycle delay of 1.0 sec to increase throughput, requiring only 1.5 min in total per experiment, for an overall acquisition time of about 5 min, including experimental setup (locking, tuning, shimming).

We also generated titration curves under high salt buffer conditions that mimic the conditions prevalent in the cell. A 10mL stock of each compound at 100uM concentration was dissolved in high salt buffer (50 mM d9-deuterated bis-trs buffer, at pH 6.5, containing 11.1 mM DSA, 200 mM NaCl, 50 mM KCl and 4 mM MgCl_2_). Compounds were divided into 12 1.5mL microcentrifuge tubes at 490 uL each and titrated with 10 uL of RNA (0.5 to 1,000 uM). The final RNA concentration in each tube increased from 0.01 uM (10 nM) to 10-20uM, while a tube with no RNA was used as control.

Titration of RNA into samples of the small molecules shown in Table 1 results in decreases in free ligand linewidth (Fig. S3), and corresponding decreases in amplitude, which are then plotted against the relevant RNA concentration (Fig. 1; Fig. S4). For proteins with compact and globular shape, these curves can be directly fit using equation 1 below, to extract binding constants ^52^. This approximation is also likely to apply to an RNA the size of the pre-miR-21 sequence used by us (Fig. S1) ^53^, whose shape approximates an ellipsoid of rotation of aspect ratio between 1.1-1.2, when hydration is considered. In the expression that follows, I_B_ is the intensity of the bound ligand peak height, I_F_ is the free ligand peak height, P_t_ is the total RNA concentration and L_t_ is the total ligand concentration. The constant c is the ratio of the bound peak width ν_B_ and the free peak width ν_F_:

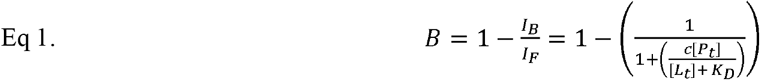

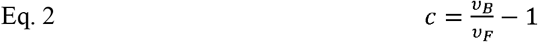

For proteins, the bound peak width (ν_B_) can be approximated by the molecular weight of the protein multiplied by a shape-related constant ρ as follows ^52^:

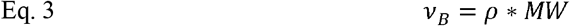

An RNA of the size of pre-miR-21 has a diameter of about 25A, accounting for hydration, and a length of 30-35A. However, the quantitative analysis of equation 1-3 might not be fully warranted when a ligand induces large changes in structure and shape upon binding; under these circumstances, variations in the line width constant c might not reliably allow measurement of bound ligand line width ν_B_, because it would change between free and bound ligand state. While the changes in hydrodynamic shape are not likely to be large given the small size of the RNA and its shape, changes in this constant would lead to uncertainties in the binding affinities.

When these conservative assumptions are made, plots of changes in ligand peak height vs added RNA concentration provide a robust semi-quantitative estimate of affinity. We then fit the data to single site binding curve models using equation 4, where B_max_ is the maximum binding capacity and represents fully titrated or broadened ligand signal, R_t_ is the total RNA concentration and NS is the slope of the nonlinear regression (non-specific binding is assumed to be linear with respect to RNA concentration):

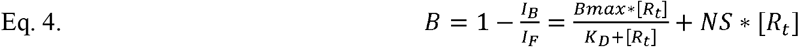

### Target-detected NMR binding analysis

The RNA sequences were dissolved in 300-500uL NMR buffer (50mM d_9_ bis-Tris pH 6.5, 50mM NaCl), heated to 95 °C for 4 min, then snap-cooled at -20 °C prior to NMR measurements. Interactions were monitored through changes in chemical shift in a series of NMR experiments, primarily 1D ^1^H and 2D ^1^H-^1^H TOCSY or ^1^H-^1^H NOESY, as needed, which were used to map the binding site of the ligand on the RNA. All pulse programs used in these experiments are standard pulse sequences provided by Bruker. Experiments in H_2_O were used to detect changes in exchangeable proton signals to assess the RNA secondary structure, while experiments in D_2_O were used to improve spectral resolution by reducing overlap of exchangeable proton signals.

After pre-miR-21 was fully titrated with a small molecule, initial RNA assignments for the complex were obtained by comparing TOCSY and NOESY spectra of the pre-miR-21:small molecule complex with those of free pre-miR-21.

#### Biochemical assays

##### RNAs

All pre-miRNA substrates were chemically synthesized by Integrated DNA Technologies (IDT) for efficient 5’-end labeling, which was done using a modified T4 PNK reaction; all sequences examined in this study are listed in Tables S1 and S2. Dicer was purchased from Creative Bio-mart (45-1 Ramsey Road, Shirley, NY 11967, USA)

RNA processing substrates were ^32^P radiolabeled using T4 polynucleotide kinase and gamma ^32^P ATP as per standard protocol. Unincorporated ATP was removed by either G25 spin columns or Zymo Research oligo clean up columns, followed by further purification by denaturing polyacrylamide gel (15 or 20%) electrophoresis.

Initial titrations were done as follows. RNA stocks in reaction buffer without DTT were folded by heating to 95°C for 2 minutes and cooling to RT. RNA was added to reaction mixtures containing either compound 52 or compound 208 at concentrations indicated in Fig. 4, and incubated for 10 minutes at 37°C, followed by enzyme addition and further incubation at 37°C for 1 hour. Aliquots of the completed reaction were run directly on a 15-20% sequencing gel for Phospho-imaging. Reaction conditions were 20 mM MES pH 6.5, 25 mM NaCl, 5 mM MgCl2, 1 mM DTT, 1% glycerol, 25 nM Dicer and approximately 50 to 100 nM RNA.

For the RNA processing experiments shown in Fig S10, the Dicer/TRBP enzyme complex was expressed and purified using published methods ^54, 55^ and stored at -80°C in 20 uL aliquots until needed. Briefly, the gene encoding human full-length Dicer was inserted into a modified pFastBac1 vector (Invitrogen), which has an N-terminal glutathione-S-transferase (GST) tag. The GST fusion proteins were expressed in HiFive insect cells following standard procedures and isolated by glutathione affinity chromatography using buffer A (20mM Tris-HCl pH 8.0, 150 mM NaCl, 2 mM DTT). After on-column cleavage by Tobacco Etch Virus (TEV) protease overnight, the proteins were eluted, concentrated, and loaded onto Superdex 200 10/300 GL column equilibrated in buffer A.

Sequences coding for human full-length TRBP2 were cloned into a modified pET28a vector with an N-terminal GB1 tag. The clones were transformed into Rosetta 2 cells and the transformants were grown in LB media at 37 °C until the cells reached a density corresponding to an OD reading of 0.8 at 600 nm. Protein expression was induced by addition of 0.2 mM IPTG at 18 °C for 20 hours. The cells were harvested, pelleted, and resuspended in 20 mM Tris-HCl pH 8.0, 500 mM NaCl, 25 mM imidazole, with 5 mM β-ME. Cells were lysed by sonication on ice then pelleted by centrifugation. The crude lysate was applied to a nickel affinity column and eluted in 20mM Tris-HCl pH 8.0, 500 mM NaCl, 250 mM imidazole buffer, with 5 mM β-ME. TEV protease was added to remove the His-GB1 tag at 4 oC overnight. The sample was concentrated and loaded onto a Superdex 200 10/300 GL column equilibrated in buffer A. The purified Dicer and TRBP2 proteins were mixed at a molar ratio of 1:1.3 and loaded onto a Superdex 200 10/300 GL column equilibrated in buffer A. The fractions containing the complex were pooled and concentrated to about 250 nM, then flash frozen for further use in processing assays.

Dicer/TRBP assays were conducted in 60 uL total volume with 25 nM RNA, 1 nM Dicer/TRBP in 1x reaction buffer (20 mM Tris at pH 7.5, 25 mM NaCl, 5 mM MgCl2, 1 mM DTT and 1% glycerol). Assays were conducted in 96-well format in a PCR thermocycler in 0.2 mL PCR well plates to maintain constant temperature and improve reproducibility. This approach allows for direct comparison of enzyme and inhibitor activity for different pre-miRNA substrates, along with parallel replicates to ensure reproducibility, especially with regards to enzyme concentration. Each reaction well contained 25 uL of RNA substrate (typical stock RNA concentration was 67 nM, but this concentration was titrated for kinetics work). Each enzyme well was filled with 50 uL of freshly diluted 2 nM Dicer-TRBP while each ligand well was filled with 50 uL of ligand at different concentrations (0-150 uM). Finally, 20 uL of reaction stop buffer (2x RNA load buffer, 95% formamide, 18 mM EDTA, 0.025% SDS, 0.1% xylene cyanol and 0.1% bromophenol blue) were added to the remaining wells.

Prior to initiating the reactions, 10 uL of ligand were added into the RNA wells using a multichannel pipette and allowed to equilibrate at 4°C for 30 min. The temperature of the thermocycler was then raised to 37°C and samples were incubated for 15 min to reach thermal equilibrium. Reactions were initiated by adding 25 uL of Dicer-TRBP to the RNA substrate wells (with or without small molecule ligand). The final enzyme concentration for all experiments was fixed to 1 nM while the final RNA and ligand concentrations were variable. The reactions were stopped by removing 5 uL from the reaction well and adding it to stop buffer wells at the indicated time points. Reactions were resolved by 20% 19:1 denaturing (8M urea) polyacrylamide gel and visualized using a Typhoon Gel imaging system (GE). Bands were quantified in ImageJ and results visualized as ratios of cleaved and uncleaved RNAs, after subtracting the background.

#### Cell-based experiments

##### Cell line maintenance

Using guidance from the Cancer Cell Line Encyclopedia (CCLE), cancer cell lines available *in house* were screened to measure levels of miR-21 expression. The human-derived cell lines AGS (gastric cancer) and AsPC-1 (pancreatic cancer) were chosen for our experiments because of the very high levels of miR-21; we hypothesized it would be difficult to regulate miR-21 in cell lines where it is not well-expressed.

Cell lines were obtained from the American Type Culture Collection and maintained per instructions from the supplier. Namely, AGS cells were cultured in F12K medium supplemented with 10% FBS, while AsPC-1 cells were cultured in RPMI, with 10% FBS added. All cells were kept at 37°C in a humidified atmosphere of 5% CO_2_ and underwent regular mycoplasma testing. All experiments were conducted when cells were 70-80% confluent.

##### Proliferation assays

The impact of the compounds on cellular proliferation was assessed with 5,000 cells in each cell line seeded in replicates of six in a 96 well plate and treated with DMSO or with compounds. After 48 hours, cells were treated with the MTS reagent and optical density was read to generate dose-response curves.

##### Quantitative real time PCR analysis

RNA was isolated utilizing Trizol (Invitrogen) and the Direct-zol RNA Miniprep kit (Zymo Research). RNA concentration was assessed with a NanoDrop ND-1000 spectrophotometer (ThermoFisher Scientific). Expression levels for all microRNAs were tested in the two representative cell lines using qRT-PCR. Cells were seeded in 6-well plates 24 hours before treatment, at a cell confluency of 50-60%. Each cell line was treated with 10 μM or 5 μM of compound or with only the DMSO solvent. RNA was collected at the 48-hour time point. Expression of miR-21 was assessed utilizing the TaqMan microRNA assay (Applied Biosystems). The complementary DNA (cDNA) was synthesized using the TaqMan Reverse Transcription Reagents kit (Applied Biosystems) and was used with the TaqMan probes and Supermix (BioRad) for qRT-PCR analysis. U6 or U48 were used as internal controls for RNA levels. All experiments were performed in triplicate, with all samples normalized to the internal controls and relative expression levels calculated using the 2^-ΔΔCt^ method.

##### Western blot analysis

Western blot analysis was conducted according to standard protocols. Briefly, proteins were collected from cells and lysed, and the Bradford assay was used to measure protein concentrations. Quantification of protein expression was conducted using the image analysis software ImageJ.

### KinomeEdge and CEREP

These experiments were conducted at Eurofin using standard protocols as reported in the company’s material; data were analyzed at Eurofin and presented in standard graphic format generated using the company software.

## ACKNOWLEDGMENTS

We wish to thank all members of the Varani group for discussion and support. The work at the University of Washington was supported by NIH MIRA grant 1 R35 GM126942 and by a gift from the Washington Research Foundation. Work at the MD Anderson was supported by grants from NIH-NCI.

## AUTHOR CONTRIBUTIONS

MDS: Experiments, data analysis and graphics, writing. BC: NMR experiments and data analysis, editing; HJZ: Cellular experiments and data analysis, writing and editing; TP: Biochemistry, editing; GLO: Data analysis, graphics, writing and editing; GAC: Experimental design, data analysis, writing, reviewing and editing; GV: Experimental design, data analysis, writing, reviewing and editing.

