## Supplementary material for "Drug-like small molecules that inhibit expression of the oncogenic microRNA-21": Suplementary figures

**Supplementary figures**

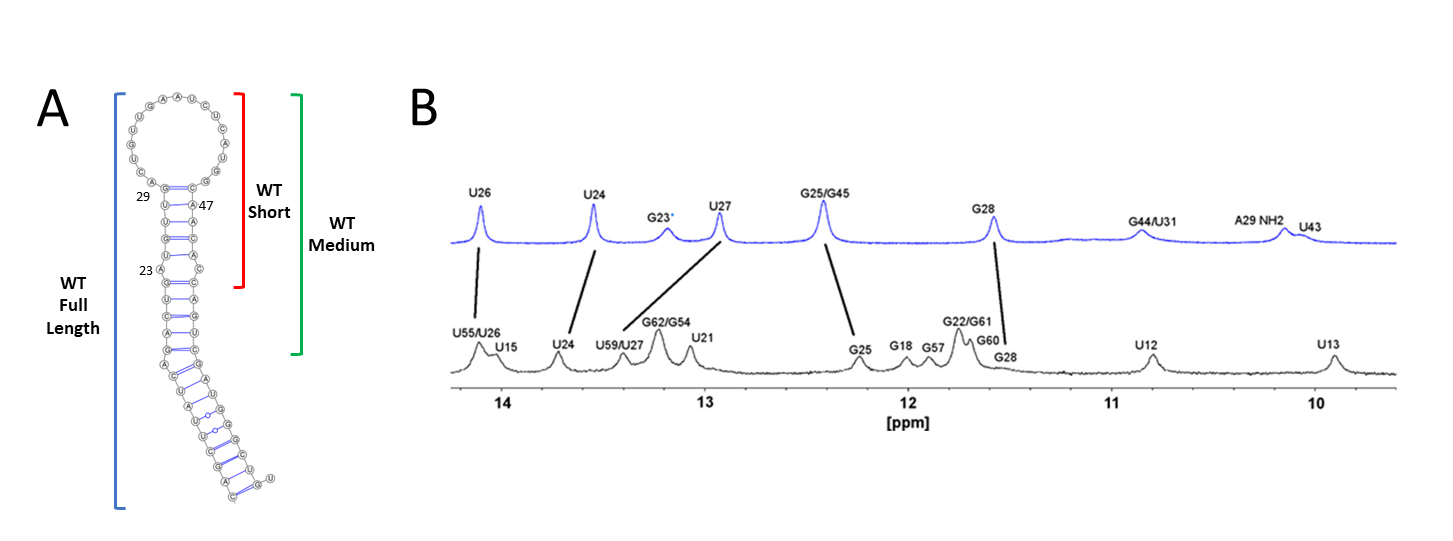

**Figure S1**. **Pre-miRNA-21 secondary structure.** (**A.)** The three miR-21 sequences used in this study, short miR-21 (nt G22-C52); medium (G18-56) and full (G8-U66) corresponding to the complete pre-miR-21 sequence. For the short RNA sequence, an A23G substitution was used to improve transcription; similarly, in the full pre-miR-21, the G8-U65 pair was swapped to CG to improve transcription. (**B**) The 1D ^1^H imino region NMR spectra for both the short transcript (blue) and the full length (bottom, black) pre-miR-21, showing the correspondence of equivalent signals (black lines), indicating identical folding; both transcripts lack NH signals from the apical loop region, which is unpaired and dynamic.

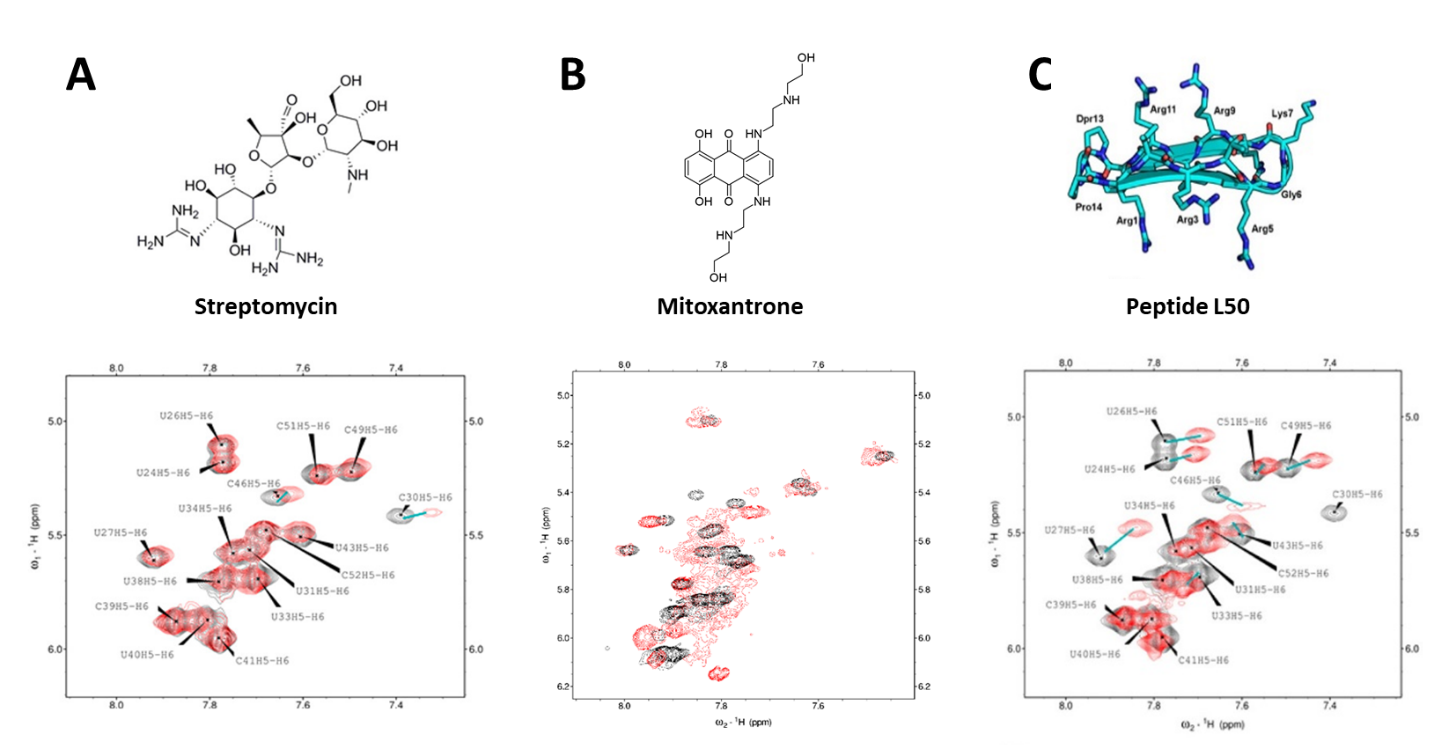

**Fig. S2** TOCSY NMR spectra illustrating responses of miR-21 to three previously proposed ligands. Signals for free pre-miR-21 are shown in black and those of the bound RNA in red. (A) Binding of the aminoglycoside streptomycin ^3^ to pre-miR-21 elicits only small changes in the spectrum; (B) mitoxantrone ^4^ binds non-specifically, broadening the NMR spectrum but without inducing specific changes indicative of a site-specific interaction. (C) L50, a peptide that binds to pre-miR-21 ^5^ induces relatively small changes in the spectra, indicating a superficial interaction with the RNA. For several other molecules we tested, we were unable to detect any interaction even at the concentrations of the NMR experiment (data not shown).

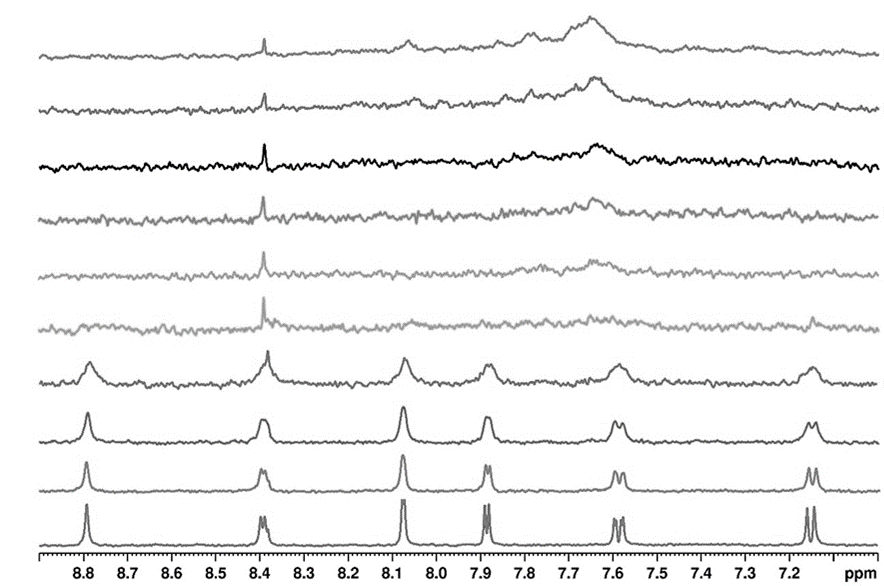

**Fig. S3A** Titration of compound 52 with increasing amounts of pre-miR-21 using a ligand-detected NMR relaxation assay demonstrates nM binding affinity (compound 52 added at 0, 0.15, 0.32, 0.65. 1.25, 2, 3, 4, 5 and 10 uM; the broad hump at 7.6 ppm is the emerging RNA signal).

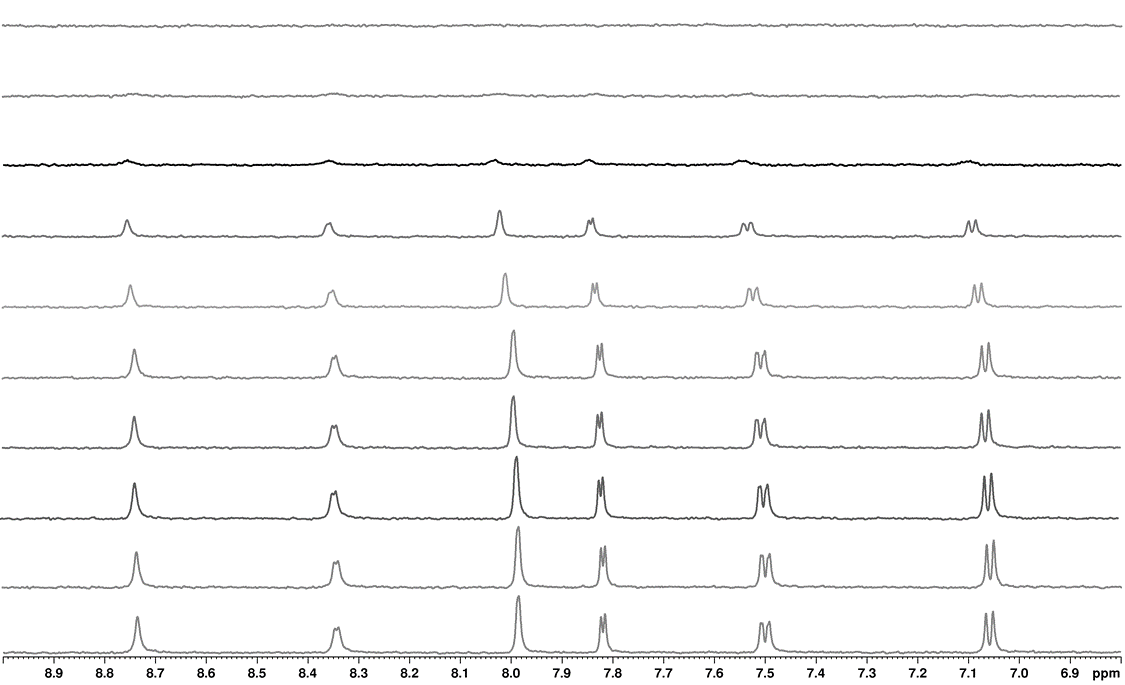

**Fig. S3B** Titration of compound 52 with increasing amounts of HIV TAR (0, 0.15, 0.32, 0.65. 1.25, 2, 3, 4, 5 and 10 uM) demonstrates weaker binding affinity, at least 10-fold less potent than binding of 52 to pre-miR-21.

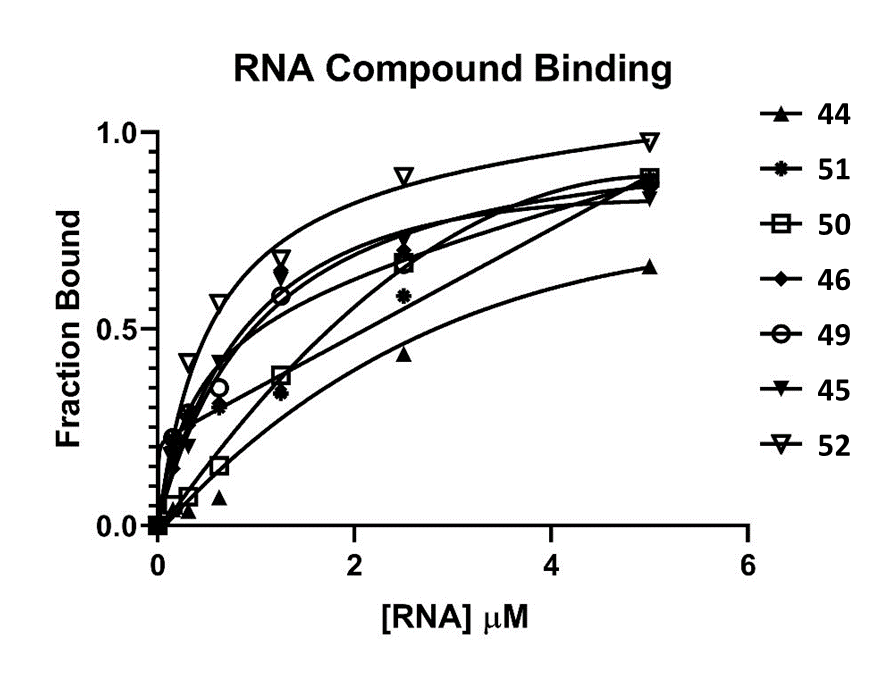

**Figure S4. Ligand-detected NMR titration of an extended set of drug-like small molecules with pre-miR-21.** A series of closely related compounds were tested for binding to pre-miR-21 using a ligand-detected 1D ^1^H NMR assay. Variants generated by moving a single nitrogen within the 2-((5-(piperazin-1-yl)pyridin-2-yl)amino)pyrido[3,4-d]pyrimidin-4(3H)-one base structure (Table 1) display significantly different affinity (5-10-fold) within the series. Titrations were performed by successive additions of RNA to a solution containing 100 μM of the relevant small molecule in a buffer containing 50 mM deuterated bis-Tris at pH 6.5, 250 mM NaCl, 50 mM KCl, and 2 mM MgCl_2_.

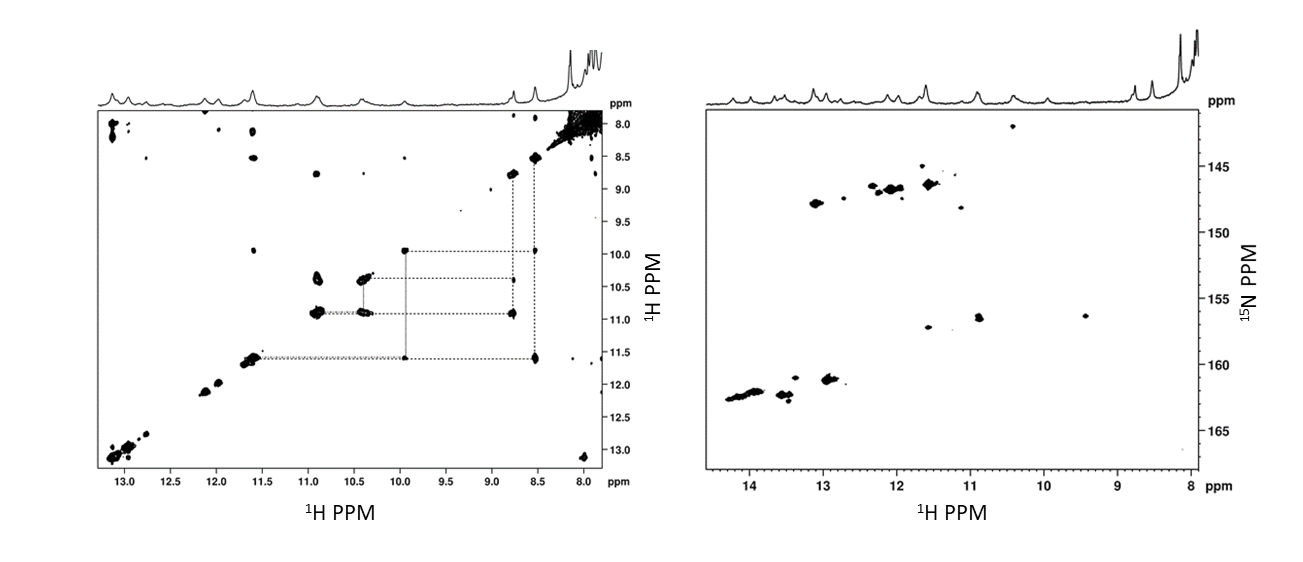

**Figure S5.** **Addition of compound 52 to pre-miR-21 results in ‘closing of the loop’ — the formation of two new GU base pairs. (A)** The NOESY imino region shows NH-NH NOEs at 10-12 ppm arising from two newly formed GU base pairs (G32:U43, and U31:G44) which appear following addition of stoichiometric amounts of compound 52 to pre-miR-21. Additional close NOE contacts to protons from the small molecule at 8.5-9 ppm are also observed, consistent with specific high-affinity binding. (B) The corresponding ^15^N HSQC provides confirmation that the NOESY signals indeed arise from GU base pairs, since the distinctive ^15^N chemical shifts identify the resonances as G’s or U’s.

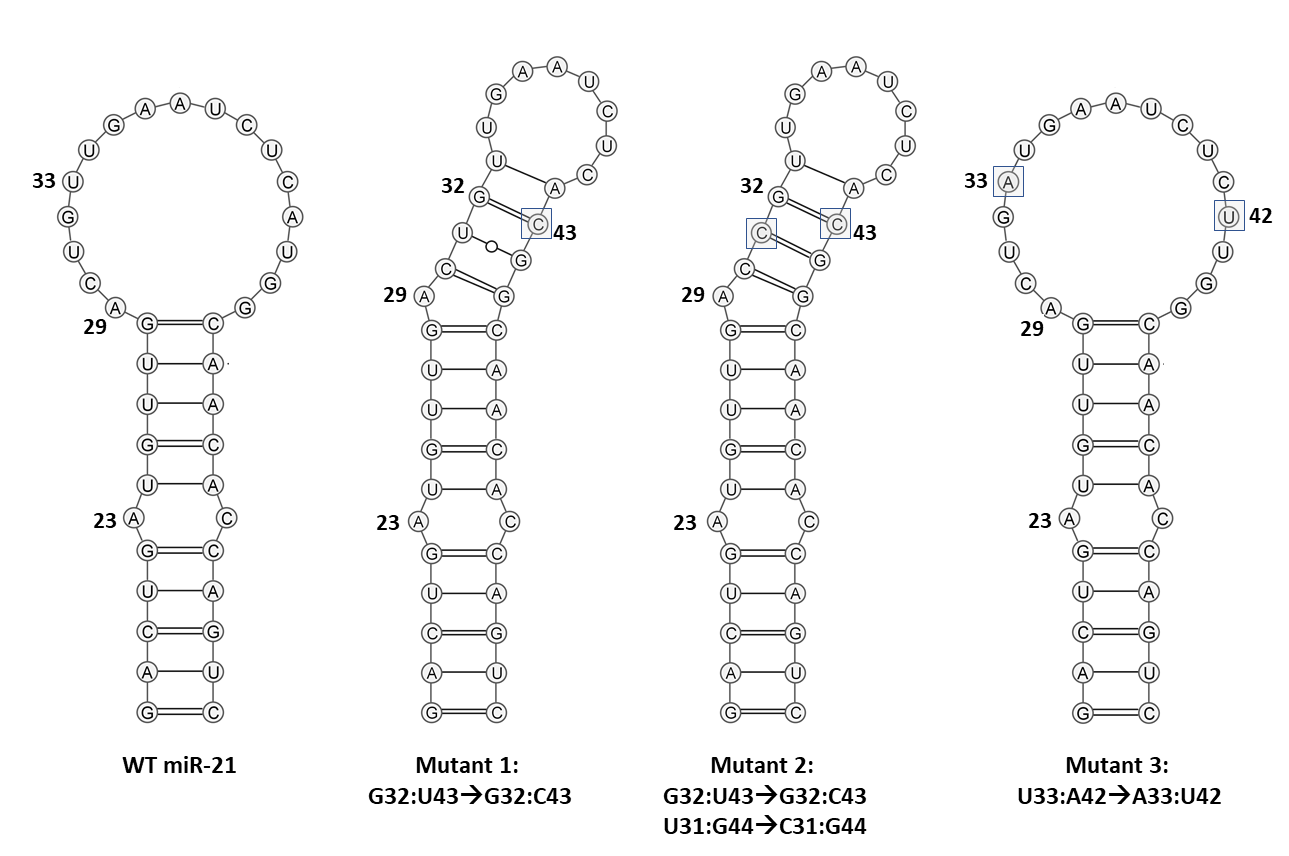

**Fig. S6A** Secondary structure of the ‘medium-length’ pre-miR-21 models used in this study, and of the 3 mutant sequences studied to control for specificity.

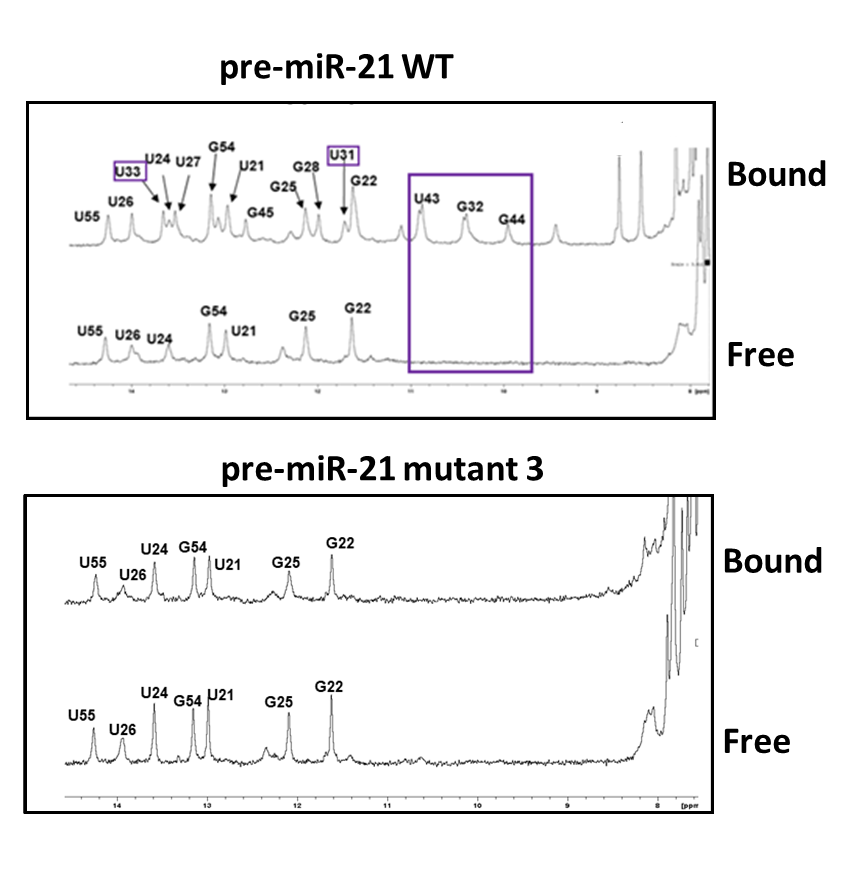

**Fig. S6B –** A single base pair inversion from UA to AU (mutant 3) prevents pre-miR-21 from adopting the ‘closed’ conformation when bound to molecule 52, essentially abrogating the stabilization of the two GU base pairs (boxed).

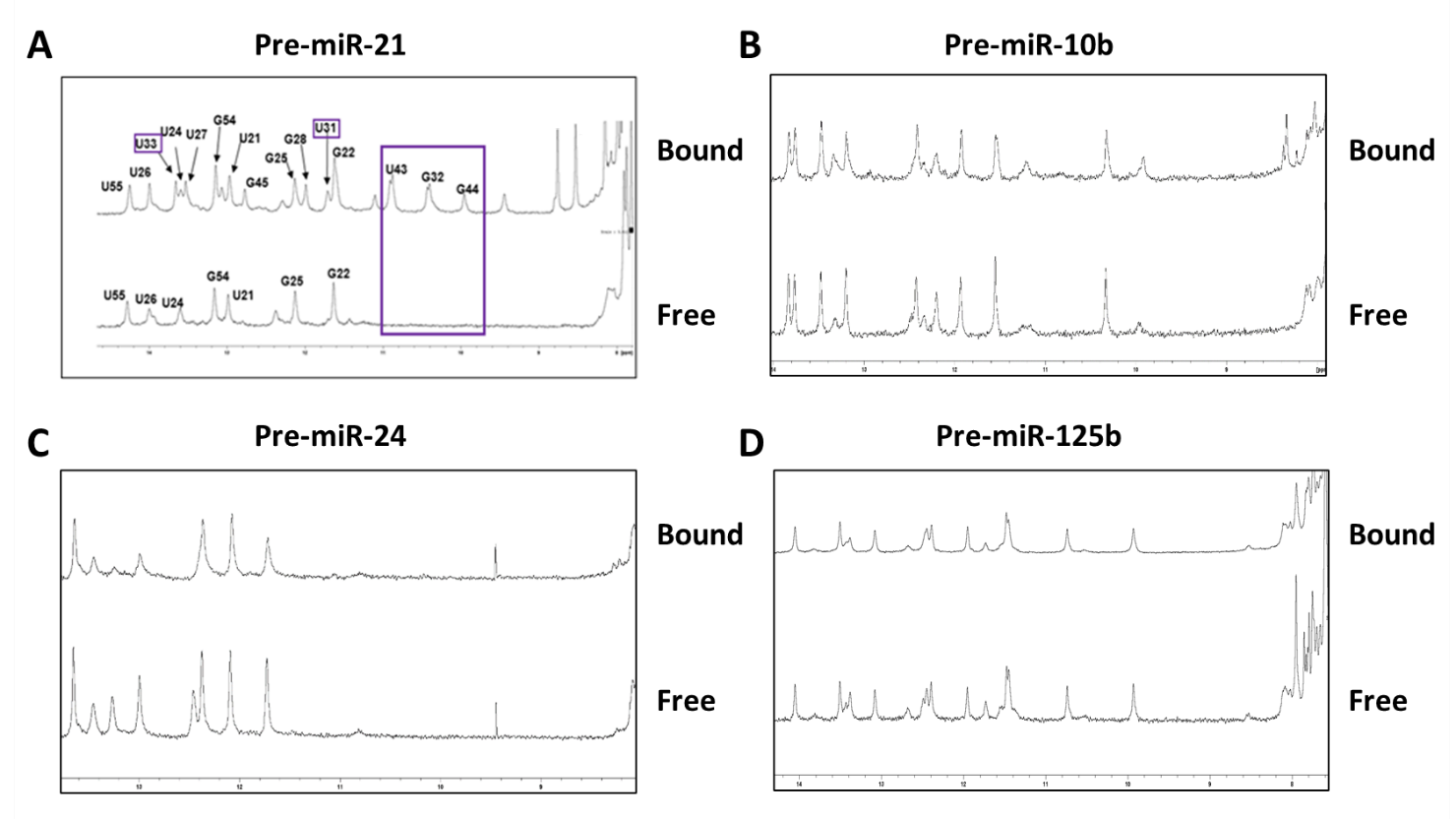

**Fig. S7 Binding of compounds 52 (left) and 45 (right) to other pre-miRNAs does not induce significant structural changes in the RNAs, in contrast to the formation of the closed-loop structure observed with pre-miR-21**. Only small changes are observed in the 1D imino NMR spectra upon binding of the small molecule to three unrelated pre-miRNAs, consistent with a superficial interaction perhaps driven by the positive charge on the piperazine ring, stacking of the heterocycle or both.

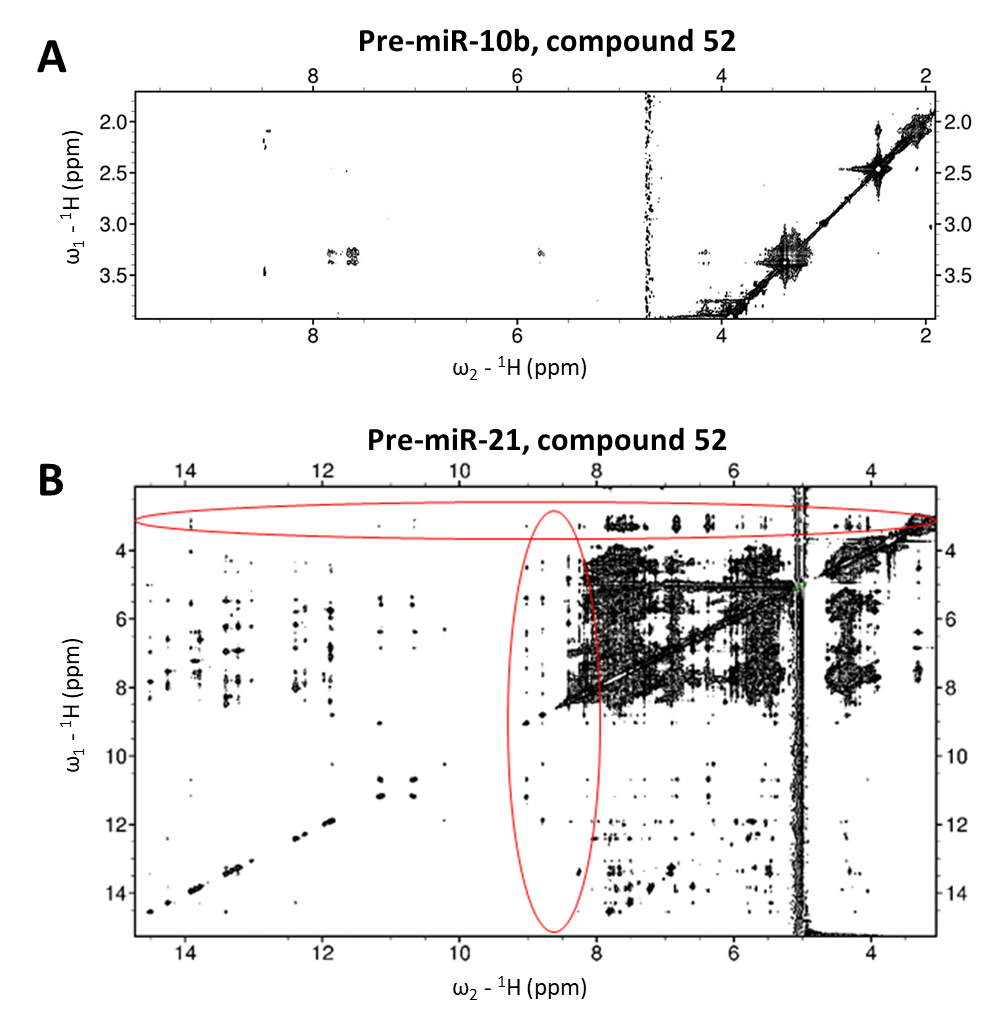

**Fig. S8 NOESY spectra illustrate the different responses of pre-miR10b and pre-miR-21 to compound 52. (A)** The interaction between molecule 52 and pre-miR-10b (top) gives rise to few intermolecular NOEs. **(B)** In contrast, the interaction between molecule 52 and pre-miR-21 gives rise to a large number of intermolecular NOEs, indicative of a stable site-specific complex. The two ovals highlight regions containing numerous intermolecular NOE interactions involving the heterocyclic ring and pyridine (vertical oval) or the piperazine (horizontal oval).

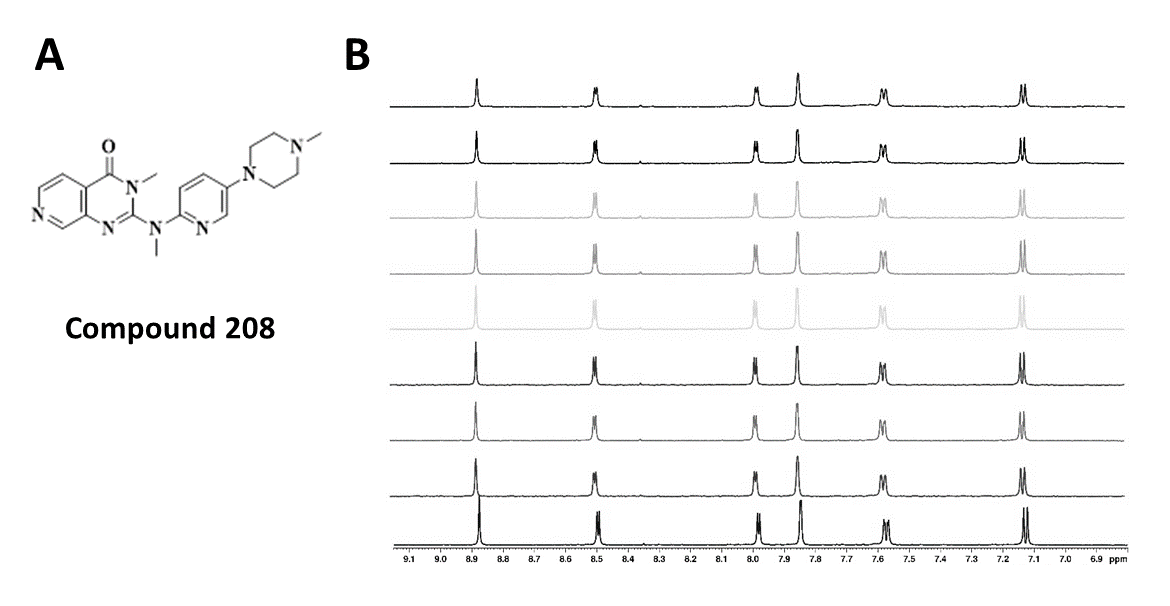

**Fig. S9** – A variant of compound 52 where each of the three NH’s is methylated does not bind to pre-miR-21; 100 uM of compound 208 was titrated with increasing amounts of pre-miR-21 (0, 0.15, 0., 0.65, 1.25, 2, 3, 4 and 5 uM). No changes in the spectra were observed, suggesting that RNA binding has been significantly reduced.

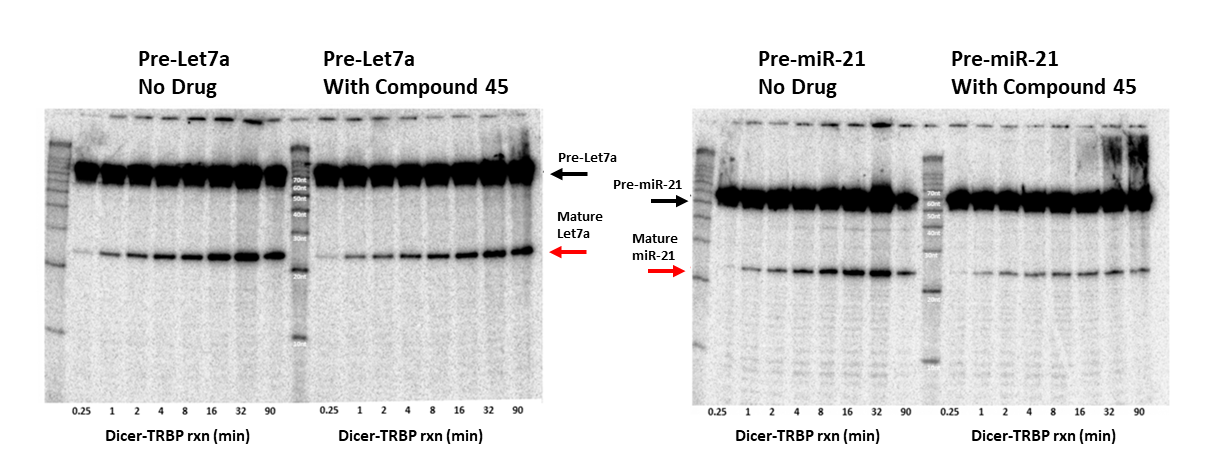

**Fig. S10. Specific inhibition of Dicer-TRBP processing by compound 45: Left**, DICER/TRBP activity on pre-Let7 with (right) or without (left) 2μM of compound 45; no significant difference in activity is observed in the presence or absence of compound 45. **Right** pre-miR-21 processing with (right) or without (left) 2μM compound 45; processing activity is visibly reduced by addition of compound 45.

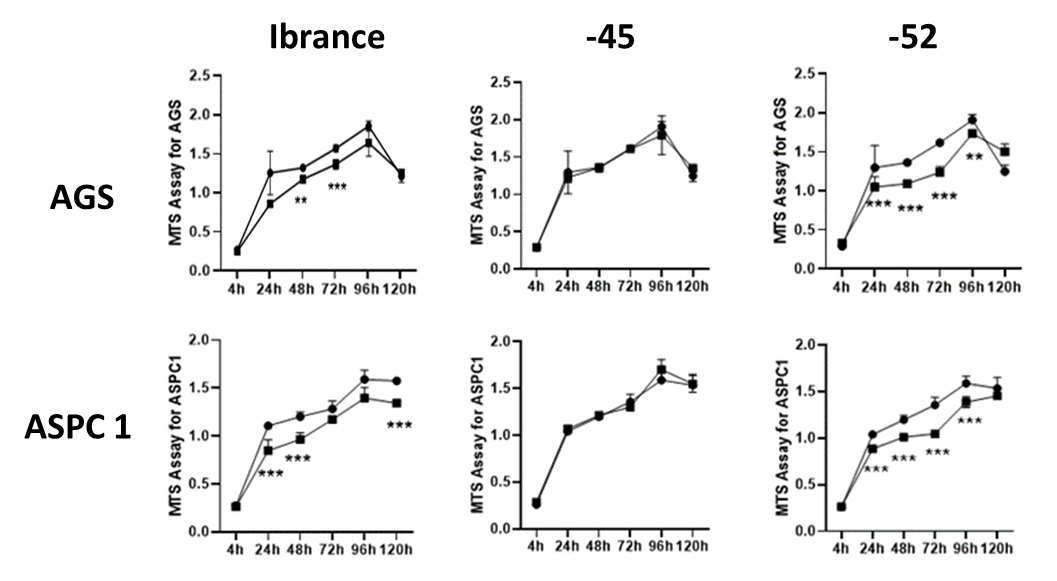

**Figure S11:** Anti-proliferative activity of 5 nmoles/1 ml, corresponding to 5 uM concentration, for compounds 45 and 52 against gastric (AGS) and pancreatic (ASPC1) cancer cell lines, measured as a function of time in a blind experimental format and compared to the kinase inhibitor Palbociclib (Ibrance). Cell viability was assayed using standard MTS assays in the indicated cell lines, following addition of each compound at 10 uM concentration (10 nmoles/1 ml); error bars show experimental uncertainty from results collected in triplicate. Palbociclib and compound 52 reduce the viability of both cell lines to a statistically significant extent (marked with asterisks) while compound 45 does not have a significant effect.

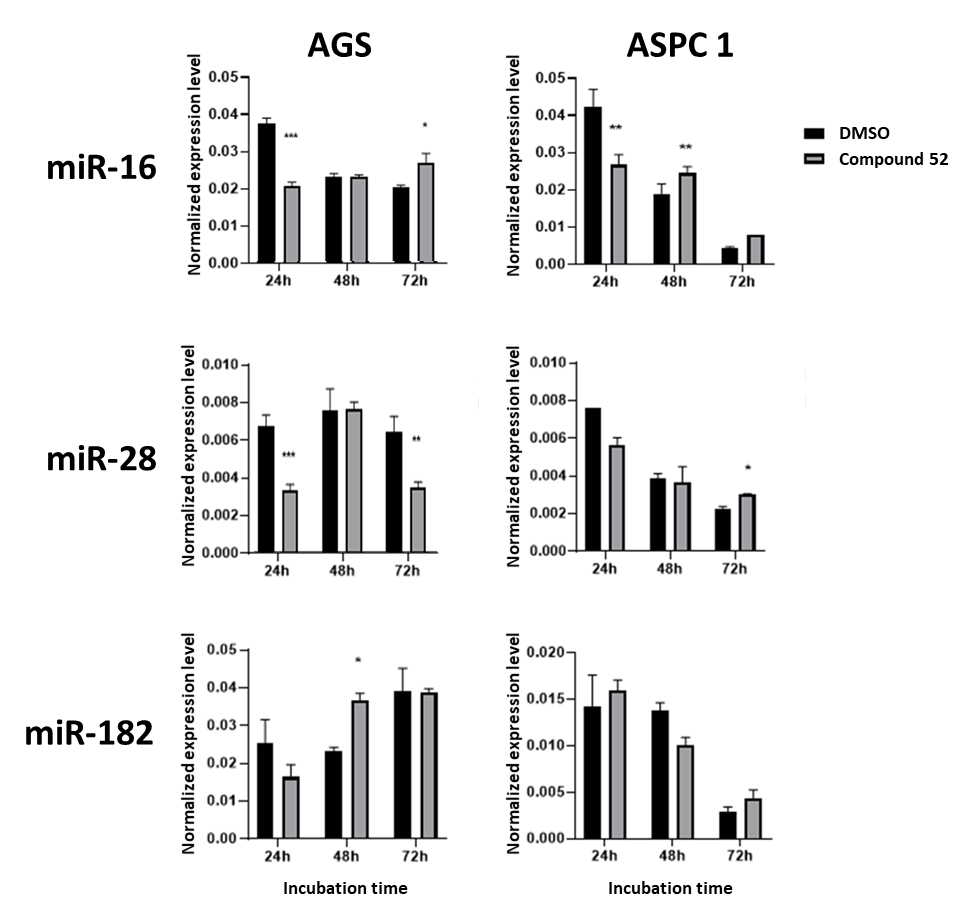

**Figure S12** The activity of compound 52 is specific for miR-21 in both pancreatic and gastric cancer cells and does not target other house-keeping miRNAs such as miR-16, miR-28 or miR-182. Levels of each microRNA were measured 48 hrs post incubation by standard qRT-PCR and normalized to internal controls (U6 snRNA).

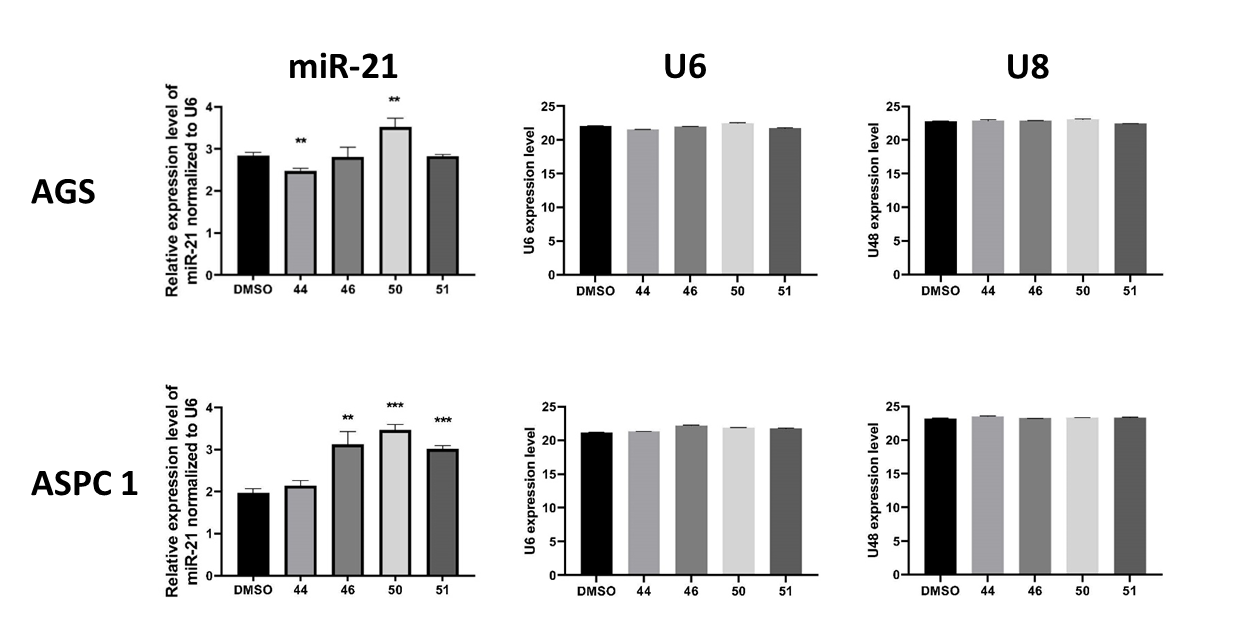

**Fig S13** Compounds 44, 46, 50 and 51 (Table 1) do not induce significant changes in miR-21 levels in either AGS or ASPC1 cancer cell lines; levels of each microRNA were measured 48 hrs post incubation by standard qRT-PCR, normalized to internal controls (U6 or U8 snRNA). The levels of U6 or U8 snRNA were not significantly affected by our compounds either.

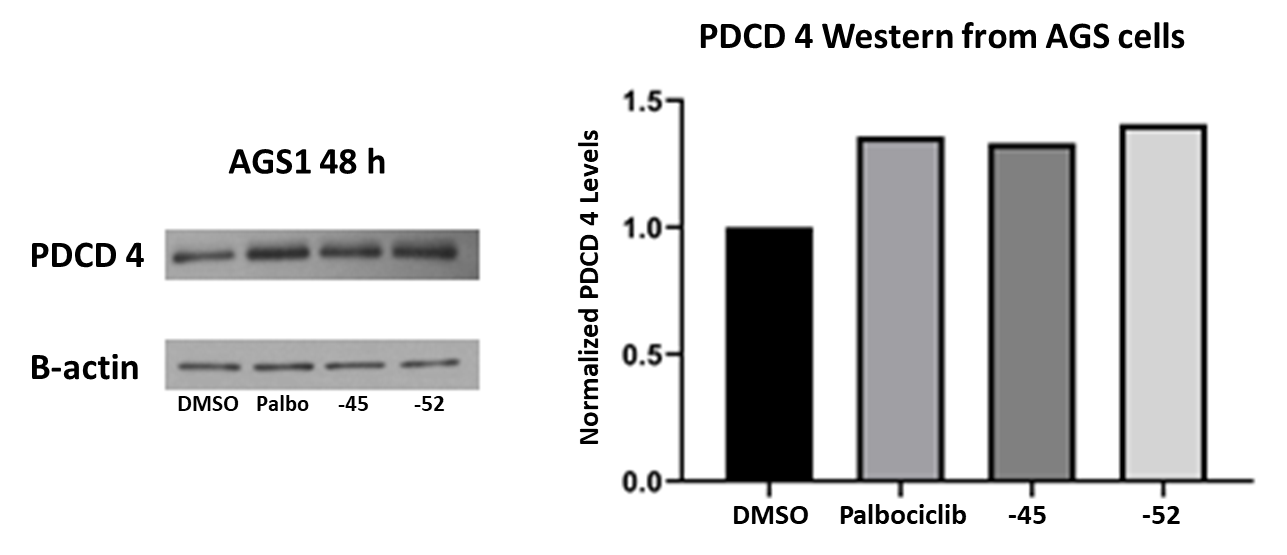

**Fig S14** Compounds 45 and 52 increase levels of PDCD4 in AGS cells following 48 hours of incubation, relative to DMSO control and normalized to β-actin levels, as measured by Western blotting (right) with an antibody specific to PDCD4. Data on the right are reported as enhancement, relative to DMSO (no compound added).

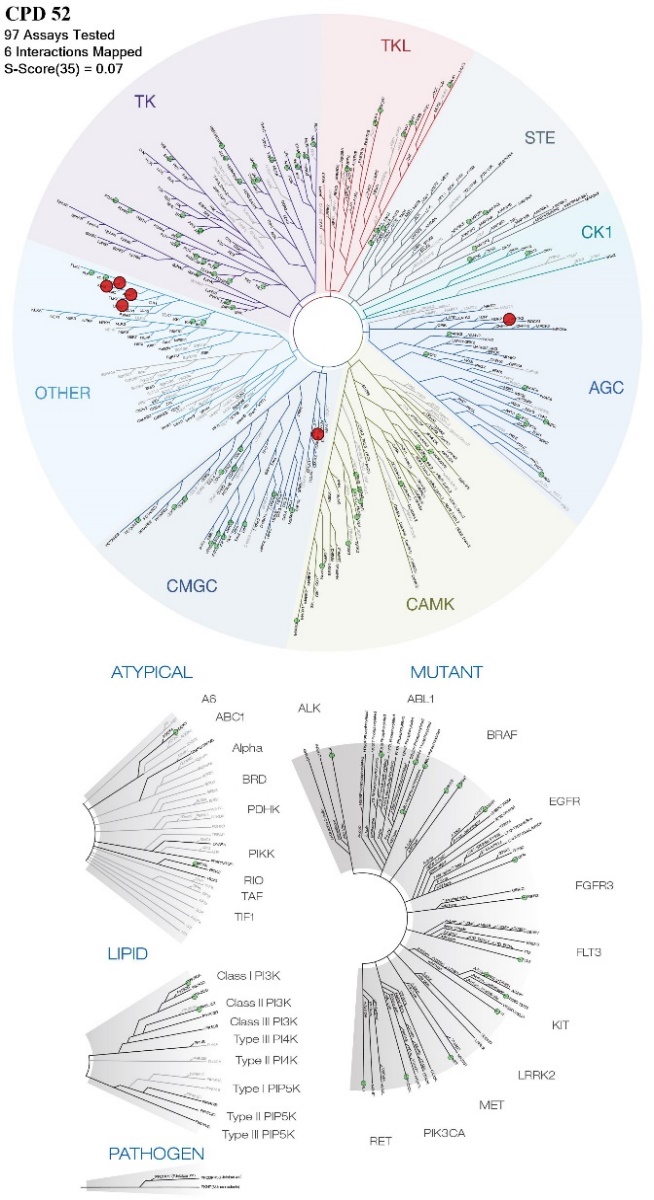

**ROCK2**

**GSK38**

**AURK (A &B)**

**PLK4 &ULK2**

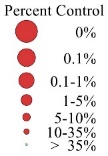

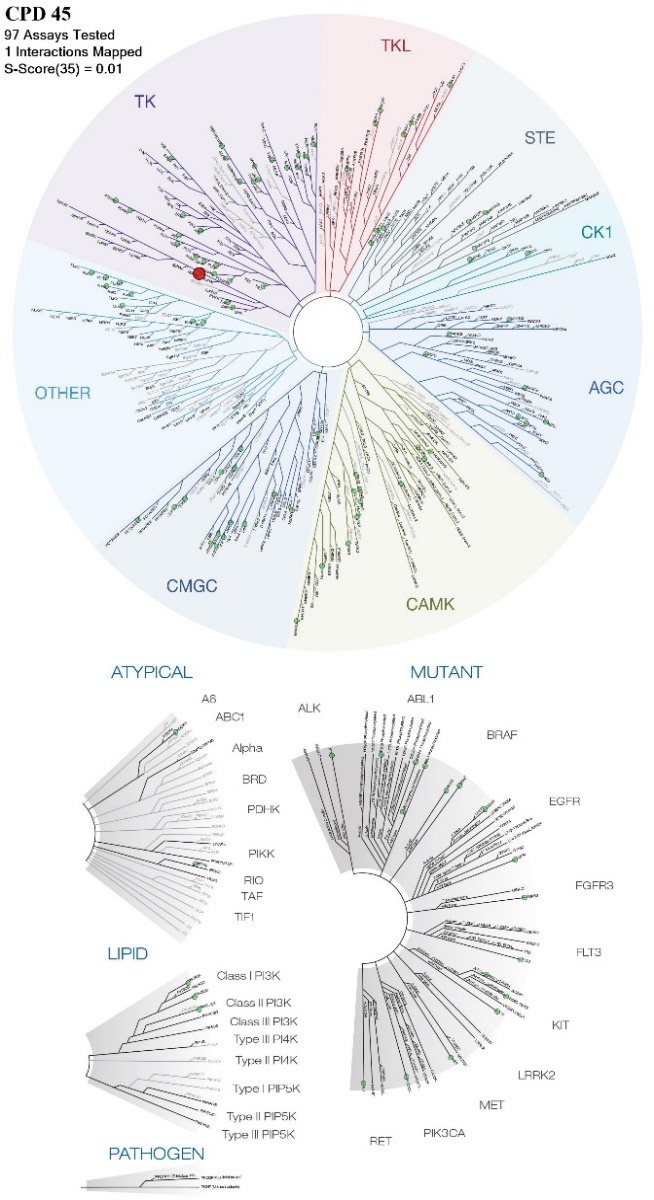

**EPHA2**

**Fig. S15** Graphical summary of a KinomeEdge assay investigating the activity of compounds 45 and 52 on approximately 100 human kinases, demonstrating only weak inhibition of a few kinases at 10uM concentration of either compound; enzymes for which inhibitory activity was observed are indicated by red circles, the size of the circle is proportional to inhibitory concentration, as indicated.

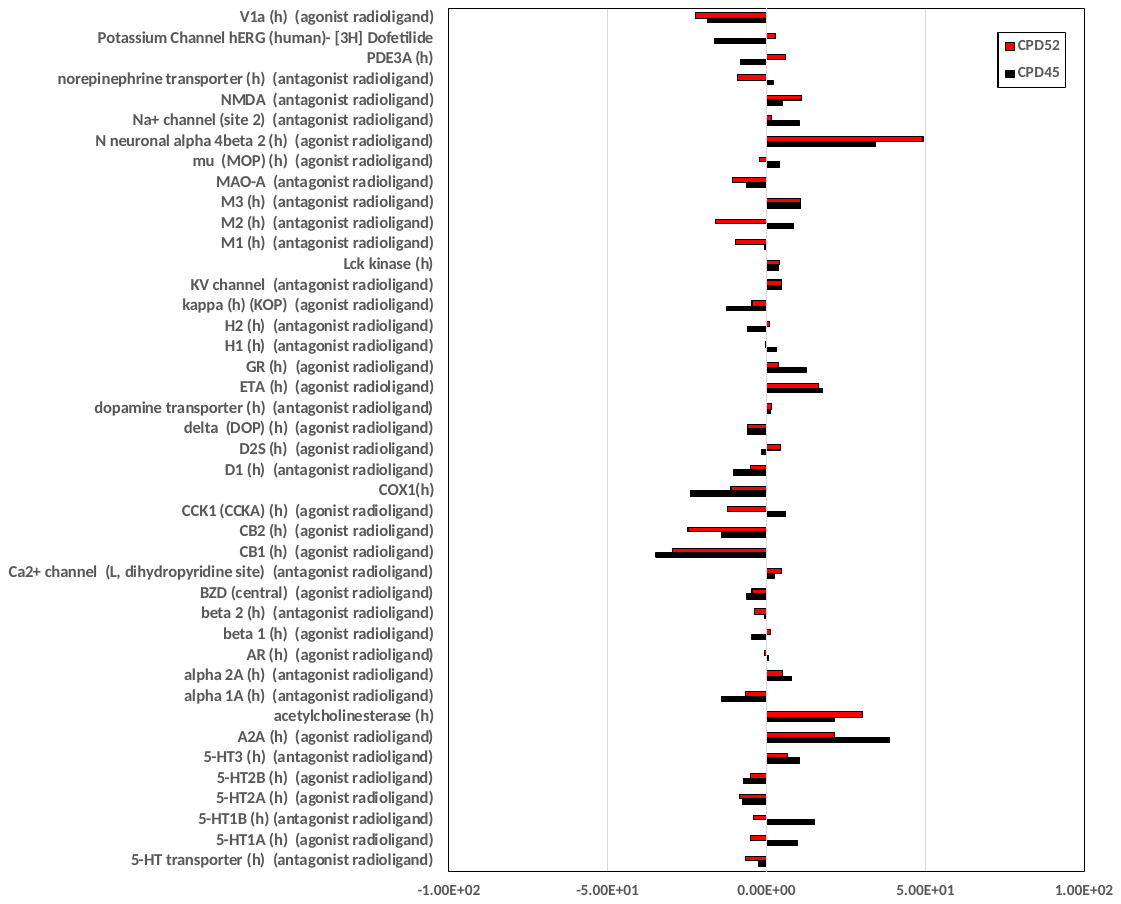

**Fig S16** Graphical summary of a CEREP assay investigating the activity of compounds 45 and 52 on a set of approximately 30 human receptors, demonstrating only weak activation on the cannabinoid receptor at 10uM concentration of compound (significant inhibition or activation is represented by bars that extend past the vertical dashed lines).

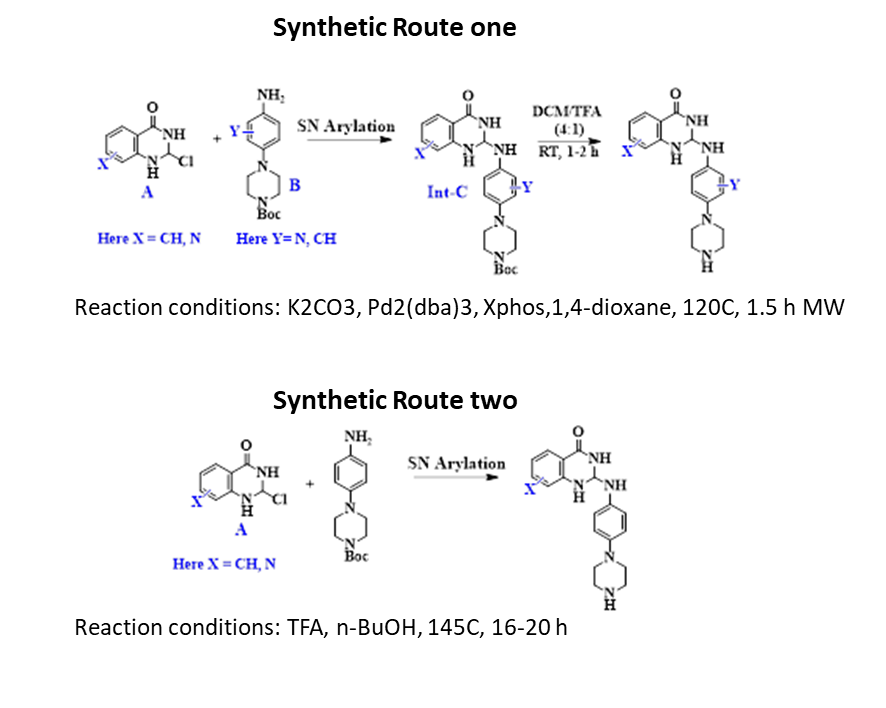

**Fig S17:** Synthetic routes used to prepare the compounds shown in Table 1.

**Table S1:** pre-miR-21 sequences and variants used in this study

| **Pre-miR-21 short (with A23G)** | GGuguug acuguugaaucucaugg caacacc |
| --- | --- |
| **Pre-miR-21 medium** | GACUGAUGUUG ACUGUUGAAUCUCAUGG CAACACCAGUC |
| **Pre-miR-21 full length (with G8:U65🡪C8:G65)** | CAGCUUAUCAGACUGAUGUUG ACUGUUGAAUCUCAUGG CAACACCAGUCGAUGGGCU**G**U |
| **WT** | GACUGAUGUUG ACUGUUGAAUCUCAUGG CAACACCAGUC |
| **Mutant 1 (U33A and A42U)** | GACUGAUGUUG ACUG**A**UGAAUCUC**U**UGG CAACACCAGUC |
| **Mutant 2 (U43C)** | GACUGAUGUUG ACUGUUGAAUCUCA**C**GG CAACACCAGUC |
| **Mutant 3 (U31C and U43C)** | GACUGAUGUUG AC**C**GUUGAAUCUCA**C**GG CAACACCAGUC |

**Table S2:** Other RNA stem-loops and miRNA hairpin sequences used for evaluating compound specificity.

| **pre-Let-7a1:** | GGGGAUGAGGUAGUAGGUUGUAUAGUUUUAGGGUCACACCCACCACUGGGAGAUAACUAUACAAUCUACUGUCUUUCCUC |
| --- | --- |
| **pre-miR-10b:** | GGGAAUUUGUGUGGUAUCCGUAUAGUCACAGAUUCCC |
| **pre-miR-24:** | GGGAGCUGAAACACAGUUGGUUUGUGUACACUGGCUCCC |
| **pre-miR-125b:** | GGGAACUUGUGAUGUUUACCGUUUAAAUCCACGGGUUCCC |
| **HIV TAR:** | GGCAGAUCUGAGCCUGGGAGCUCUCUGCC |
